# Pharmacological Characterization of SDX-7320/Evexomostat: a Novel Methionine Aminopeptidase Type 2 Inhibitor with Anti-Tumor and Anti-Metastatic Activity

**DOI:** 10.1101/2024.02.29.582701

**Authors:** Peter Cornelius, Benjamin A. Mayes, John S. Petersen, David J. Turnquist, Pierre J. Dufour, James M. Shanahan, Bradley J. Carver

## Abstract

Methionine aminopeptidase type 2 (MetAP2) is a ubiquitous, evolutionarily conserved metalloprotease fundamental to protein biosynthesis which catalyzes removal of the N-terminal methionine residue from nascent polypeptides. MetAP2 is an attractive target for cancer therapeutics based upon its over-expression in multiple human cancers, the importance of MetAP2-specific substrates whose biological activity may be altered following MetAP2 inhibition, and additionally, that MetAP2 was identified as the target for the anti-angiogenic natural product, fumagillin. Irreversible inhibition of MetAP2 using fumagillin analogs has established the anti-angiogenic and anti-tumor characteristics of these derivatives, however, their full clinical potential has not been realized due to a combination of poor drug-like properties and dose-limiting CNS toxicity. This report describes the physicochemical and pharmacological characterization of SDX-7320 (evexomostat), a polymer-drug conjugate of the novel MetAP2 inhibitor (MetAP2i) SDX-7539. *In vitro* binding, enzyme and cell-based assays demonstrated that SDX-7539 is a potent and selective MetAP2 inhibitor. In utilizing a high molecular weight, water-soluble polymer to conjugate the novel fumagillol-derived, cathepsin-released, MetAP2i SDX-7539, limitations observed with prior generation, small-molecule fumagillol derivatives were ameliorated including reduced CNS exposure of the MetAP2i, and prolonged half-life enabling convenient administration. Multiple xenograft and syngeneic cancer models were utilized to demonstrate the anti-tumor and anti-metastatic profile of SDX-7320. Unlike polymer-drug conjugates in general, reductions in small molecule-equivalent efficacious doses following polymer conjugation were observed. SDX-7320 has completed a Phase 1 clinical safety study in late-stage cancer patients and is currently being evaluated in multiple Phase 1b/2 clinical studies in patients with advanced solid tumors.

## Introduction

The cytoplasmic metalloproteases methionine aminopeptidase types 1 and 2 (MetAP1 and MetAP2) are fundamental in cellular function across prokaryotes and eukaryotes, primarily responsible for N-terminal methionine cleavage of proteins during *de novo* biosynthesis (1). A limited number of MetAP2-selective substrates have been reported to date: thioredoxin-1, cyclophilin A, eukaryotic elongation factor 2, Rab-37, glyceraldehyde 3-phosphate dehydrogenase, 14-3-3γ and SH3 binding glutamic acid rich-like protein (2–4). Inhibiting the aminopeptidase activity of MetAP2 causes these substrates to retain their initiator methionine, thereby affecting post-translational modifications, protein folding, cellular localization, and stabilization (2,3,5,6).

Fumagillin is a natural product isolated from the fungus *Aspergillus fumigatus Fresenius* which has been shown to irreversibly bind selectively to MetAP2 (but not MetAP1) and inactivate its aminopeptidase activity (1). Fumagillin-bound MetAP2 is stabilized and accumulates intracellularly, exerting additional pharmacodynamic effects including inhibition of MAPK activity (7,8). Therefore, inhibition of MetAP2 elicits multiple downstream effects leading to various physiological responses (9–11) including inhibition of tumor growth (12), metastasis (13), and angiogenesis, due to endothelial cell cycle arrest in the late G1 phase (14,15). MetAP2 has been reported to be overexpressed in many malignancies including breast, prostate, colorectal, esophageal, lung, lymphomas mesothelioma, and cholangiocarcinoma (16–20).

Multiple fumagillin-based MetAP2 inhibitors have been evaluated clinically as anti-cancer compounds. TNP-470 is a fumagillin derivative with 50-fold higher potency *in vitro* than the natural product, for which synergistic activity was observed in preclinical studies when combined with cytotoxic chemotherapeutics (21–23). Despite promising clinical efficacy, TNP-470 suffered from dose-limiting neurological side effects, presumably due to penetration of the small molecule across the blood-brain barrier (24). Other challenges to its clinical development included poor aqueous solubility and a pharmacokinetic (PK) profile which necessitated frequent intravenous (IV) dosing due to its short half-life (9,25).

Polymer-drug conjugation is a well-documented approach that presents potential benefits with respect to the limitations associated with drugs like TNP-470 (26). As a hydrophilic polyacetal linked to a fumagillol derivative, the polymer-drug conjugate (PDC) XMT-1107 reached Phase 1 clinical studies in late-stage cancer patients (27). However, high doses were required to elicit pharmacodynamic effects and development was halted (28). A fumagillol analog derived from TNP-470, conjugated to a hydroxpropylmethacrylamide (HPMA) copolymer backbone displayed favorable preclinical characteristics in both neurobehavioral tests as well as tumor models, however it did not undergo clinical evaluation (29). Importantly, poly(HPMA) with other pendant small molecules has not been found to induce signs of toxicity nor immunogenicity (e.g. PK1, FCE28068 (30)). Alternate non-fumagillol derived PDCs using the HPMA copolymer backbone have been investigated clinically at average daily doses >1.5 g/m^2^ (e.g., AP5280; (31)). With respect to these PDCs, total drug doses in excess of 1.0 g/m^2^/day were typically required for clinical anti-tumor efficacy.

Based on the encouraging anti-tumor efficacy profile of TNP-470 and other fumagillin-based inhibitors of MetAP2, a novel fumagillol-derived small molecule was sought which when conjugated to a high molecular weight polymeric system would yield a prodrug suitable for clinical development. This report describes the structure, synthesis and pharmacological characterization of SDX-7320 (INN: evexomostat), a PDC of the novel, selective MetAP2 inhibitor SDX-7539, which displays anti-tumor and anti-metastatic properties with an improved safety profile versus previous small molecule fumagillin analogs.

## Materials and Methods

### Synthesis of SDX-7320/evexomostat

The chemical structure of SDX-7320 is shown in Figure 1, along with the pharmacologically active small molecule MetAP2 inhibitor SDX-7539. The preparation and characterization of SDX-7320 and SDX-7539 are described in the Supplementary Data, along with the related compounds SDX-9246, SDX-9280, SDX-9178, TNP-470 and CKD-732. The benzenesulfonic acid salt form of SDX-7539 (SDX-9402) was used for *in vitro* experiments.

**Figure 1.**
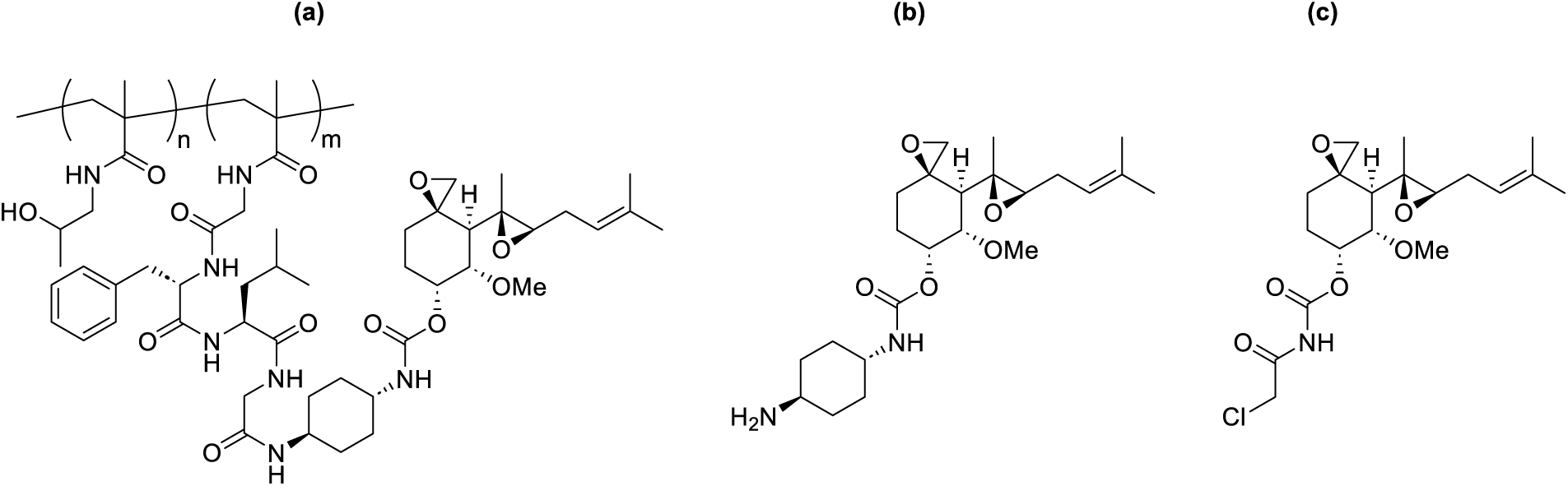
Structures of (a) polymer-drug conjugate SDX-7320 (n ∼ 90 mol%; m ∼ 10 mol%), and small-molecule MetAP2 inhibitors (b) SDX-7539 and (c) TNP-470.

### Enzyme-mediated metabolism of SDX-7320 *in vitro*

Cathepsin isoforms were purchased from the suppliers shown in the Supplementary Data. Cathepsin isoforms and human neutrophil elastase (HNE) were tested for their ability to catalyze the release of SDX-7539 from SDX-7320 *in vitro*, using manufacturer-recommended assay conditions, as described in the Supplementary Data.

### MetAP1 and MetAP2 enzyme activity assays

Enzyme activity was measured at room temperature using a coupled assay system (R&D Systems) containing the fluorogenic substrates Met-Gly-Pro-AMC (for MetAP1) or Met-AMC (for MetAP2). Reactions for MetAP2 contained 10 μg/mL hMetAP2, 5 μg/mL DPPIV, and 1 μM Met-AMC, while MetAP1 reactions contained 4 μg/mL MetAP1, 1 μg/mL DPPIV, and 1 μM Met-Gly-Pro-AMC in 50 mM HEPES, 0.1 mM CoCl_2_, 100 mM NaCl, pH 7.5. Test compounds were diluted in DMSO, then diluted in 96-well plates containing enzymes, followed by addition of the fluorogenic substrate. Fluorescence was measured over time using excitation and emission wavelengths of 350 nm and 440 nm.

### Expression of MetAP2 and MetAP2-SDX-7539 co-crystal structure

Details of the expression of MetAP2, co-crystallization with SDX-7539, data reduction and refinement are provided in the Supplementary Data. The final refined structure was deposited in the PDB under code 8OXG.

### Human Umbilical Vein Endothelial Cell (HUVEC) proliferation

HUVECs (from the American Type Culture Collection, ATCC) were cultured in EBM-2 medium with EGM-2 (Lonza #CC-3156 and CC4176, respectively) containing 2% FBS and 2X gentamicin. Cells were plated into 96-well plates (5,000 cells/well) and incubated overnight. Test compounds were initially diluted in DMSO, then diluted into culture medium listed above containing 2%FBS, 2X gentamicin (final DMSO = 0.1%). After 72 hours, 37°C Promega Substrate CellTiter 96 Aqueous One Solution Reagent was added to each well. After 3 hours at 37°C, the OD at 490 nm was measured, then corrected for baseline by subtracting the average A_490_ of the wells with no compound added. Inhibitory potency was calculated using GraphPad’s non-linear regression analysis (“log(inhibitor) vs response - Variable slope (four parameters)”).

### In Vivo Studies

All animal procedures were carried out under the institutional guidelines of each organization and were approved by their respective Institutional Animal Care and Use Committees.

- Rat Functional Observation Battery (FOB) Evaluation of the neurobehavioral safety of SDX-7320 in male experimentally naïve CD® [Crl:CD®(SD)] rats (approximately 8 weeks of age at the start of the study) was conducted under good laboratory practice (GLP) conditions following single subcutaneous doses (ten/group) of 3, 10 or 30 mg/kg. Assessments of neurobehavioral effects and general toxicity were based on mortality and FOB evaluations (32), clinical observations, and body weights. FOB evaluations were conducted pre-dose (Day −1) and at 12- and 36-hours post-dose. A detailed description of the FOB and the results can be found in the Supplementary Data.
- A549 Non-small Cell Lung Cancer (NSCLC) Xenograft Model The A549 human NSCLC cell line was obtained from ATCC. Female Athymic Nude mice (Hsd:Athymic Nude-Fox-n1^nu^ from Harlan), were inoculated subcutaneously in the right hind flank with 0.1 mL of a 50% RPMI-1640 / 50% Matrigel™ (BD Biosciences) mixture containing a suspension of A549 tumor cells (approximately 5×10^6^ cells/mouse). At time of inoculation, the mice were 5-6 weeks old. Twelve days following inoculation, tumors were measured using a digital caliper. Sixty mice with tumor volumes of 107-180 mm^3^ (mean tumor volume of 142 mm^3^) were randomized into groups of ten mice/group. Tumor volume and body weight were recorded twice weekly. All test agents were dosed IV (tail vein). SDX-7320 was dosed every four days (Q4Dx7) while TNP-470 and SDX-7539 were dosed every other day (QODx10). All mice were sacrificed twenty-nine days after initiating dosing.
- EO771 Triple Negative Breast Cancer (TNBC) Allograft Model Female C57Bl/6j mice that had undergone surgical ovariectomy at 4 weeks of age to mimic the postmenopausal state were obtained from Jackson Laboratory. EO771 tumor cells (a model of TNBC, obtained from CH3 Biosystems) were implanted under ether anesthesia into the fourth mammary fat pad when mice were approximately 18 weeks of age. Treatment with vehicle (5% mannitol/water, subcutaneous (SC), Q4D) or SDX-7320 (24 mg/kg, SC, Q4Dx4) was initiated when tumors were approximately 20 mm^3^. All mice were euthanized on Day 15 when tumor and other tissues were excised, and a portion of tumor tissue was fixed in buffered formalin and processed for histology and immunohistochemistry (IHC; see Supplemental Data for IHC details).
- MDA-MB-231 TNBC Model in Chick Embryos MDA-MB-231 cells were obtained from ATCC. The effect of SDX-7320 on tumor growth, angiogenesis and metastatic invasion was evaluated in tumors grown in fertilized White Leghorn eggs as described in the Supplementary Data.
- B16-F10 Syngeneic Lung Metastasis Model Female C57/Bl6 mice (CrTac:C57BL/6NTac) were obtained from Taconic. The B16-F10 mouse melanoma tumor cell line was from ATCC. Cultures were maintained in DMEM (Gibco, Invitrogen), supplemented with 5% fetal bovine serum, and housed at 37°C in a humidified 5% CO_2_ atmosphere. Beginning 24 hours after tumor cells were injected *via* tail vein, drug treatments were administered IV as follows (eight mice/group): vehicle (saline), TNP-470 (30 mg/kg), and SDX-7320 (25 mg/kg). Vehicle and SDX-7320 were dosed every fourth day starting on Day 1 for three treatments (Q4Dx3) while TNP-470 was dosed every other day for six treatments (Q2Dx6). All animals were euthanized on Day 15 and lungs were excised. Excised lungs were placed under a stereomicroscope and visible B16-F10 lung metastases were counted and recorded. Statistical analysis was performed with GraphPad Prism® v6.0 software. Differences in lung metastasis counts were confirmed using a Student T-test (two-tailed, two-sample equal variance).
- BT-474 HER-2+, estrogen receptor (ER)+ BC Xenograft Model Female Athymic Nude mice were obtained from Charles River and implanted with subcutaneous, slow-release estrogen (90-day, 0.72 mg β-estradiol; Innovative Research of America). Three days after receiving the estrogen pellets, mice were injected in the fourth mammary fat pad with 2×10^7^ BT-474 cells (a model of hormone receptor+/HER-2+ breast cancer), suspended in 50% Matrigel / 50% RPMI-1640). When tumors exceeded 95 mm^3^, they were eligible for randomization (eight mice/group). SDX-7320 was prepared fresh for each dose (administered SC/Q4Dx9) in 5% mannitol/water. At necropsy, tumors were excised, and a portion snap-frozen and stored at −80°C until shipped for RNA sequencing.

### RNA isolation, sequencing and analysis

Total RNA from BT-474 tumors was extracted using Qiagen RNeasy Plus Universal mini kit following manufacturer’s instructions (Qiagen). The RNA samples were quantified using Qubit 2.0 Fluorometer (Life Technologies) and RNA integrity was checked using TapeStation (Agilent Technologies). The RNA sequencing library was prepared using the NEBNext Ultra RNA Library Prep Kit for Illumina using manufacturer’s instructions (New England Biolabs), validated on TapeStation (Agilent Technologies), and quantified by Qubit 2.0 Fluorometer (Invitrogen) and quantitative PCR (KAPA Biosystems). The sequencing library was clustered on one lane of a flowcell and loaded on the Illumina HiSeq instrument (4000 or equivalent) according to manufacturer’s instructions. The sample was sequenced using a 2×150bp Paired End (PE) configuration. Image analysis and base calling were conducted by HiSeq Control Software (HCS). Raw sequence data (.bcl files) generated were converted into fastq files and de-multiplexed using Illumina’s bcl2fastq 2.17 software. One mismatch was allowed for index sequence identification. Analysis was carried out in the Watershed® Cloud Data Lab (CDL). Raw RNA-sequence reads were aligned to the human reference genome (GRCh37/hg19) with NCBI RefSeq annotations using STAR v2.7.5c. The RNASeq data were uploaded to the GEO database (accession number GSE230687). Principal component analysis (PCA) was performed using the PCA function within the python package scikit-learn. Normalized gene expression was used as input to find the first 10 components of global gene expression.

### Data availability statement

Data for this study were generated at multiple contract research organizations. Reports containing raw data as well as derived data supporting the findings of this study are available from the corresponding author upon reasonable request. X-ray crystallography data is available from the Protein Data Bank (PDB) under code 8OXG. Gene expression data is available in the GEO database under accession number GSE230687.

## Results

### Preparation and characterization of the PDC SDX-7320

The polymer drug conjugate SDX-7320 features a poly(methacrylamide) backbone functionalized with 2-hydroxypropyl groups and a glycyl-L-phenylalanyl-L-leucyl-glycyl-(GFLG) linker bearing the novel fumagillin-derived MetAP2 inhibitor SDX-7539 (Figure 1). The small molecule SDX-7539 is released from the tetrapeptide linker by enzymatic cleavage (see below). After conjugation of SDX-7539 to the copolymer *via* aminolysis of an active ester, the resultant PDC was purified by ultrafiltration and lyophilized to give SDX-7320 as an amorphous powder on 100-gram scale. Unconjugated SDX-7539 was not detected by HPLC analysis (LOD 0.02%). Small molecule SDX-7539 loading was measured to be 19% by weight using HPLC. SDX-7320 average molecular weight (M_w_) was determined by size exclusion chromatography with refractive index detection to be 29 kDa (PDI 1.49) relative to a calibration curve established using poly(methyl methacrylate) standards. Aqueous solubility of SDX-7320 was greater than 200 mg/mL (25°C). SDX-7320 was observed to have a monomodal particle size distribution with Z-average 10 nm (PDI 0.23) as measured by dynamic light scattering in 5% aqueous mannitol.

### Enzyme-mediated metabolism of SDX-7320 *in vitro*

The tetrapeptide linker “GFLG” has been previously used to create PDCs and is a substrate for lysosomal proteases such as cathepsin B (CTSB; 47). Using a standard set of conditions, multiple cathepsin isoforms were profiled for their ability to catalyze the release of SDX-7539 from SDX-7320 *in vitro* (Table S2). The results showed that several cathepsin isoforms were capable of releasing SDX-7539 from SDX-7320, and Cathepsin S (CTSS) was the most active isoform against SDX-7320. Additional cathepsin isoforms with catalytic activity against SDX-7320 included L, H, C, K and B (Figure 2). However, differences in each enzymes’ pH optima, requirement for pre-activation with CTSL as well as specific activity limit the direct comparison of catalytic efficiency between the enzymes for cleavage of SDX-7320.

**Figure 2.**
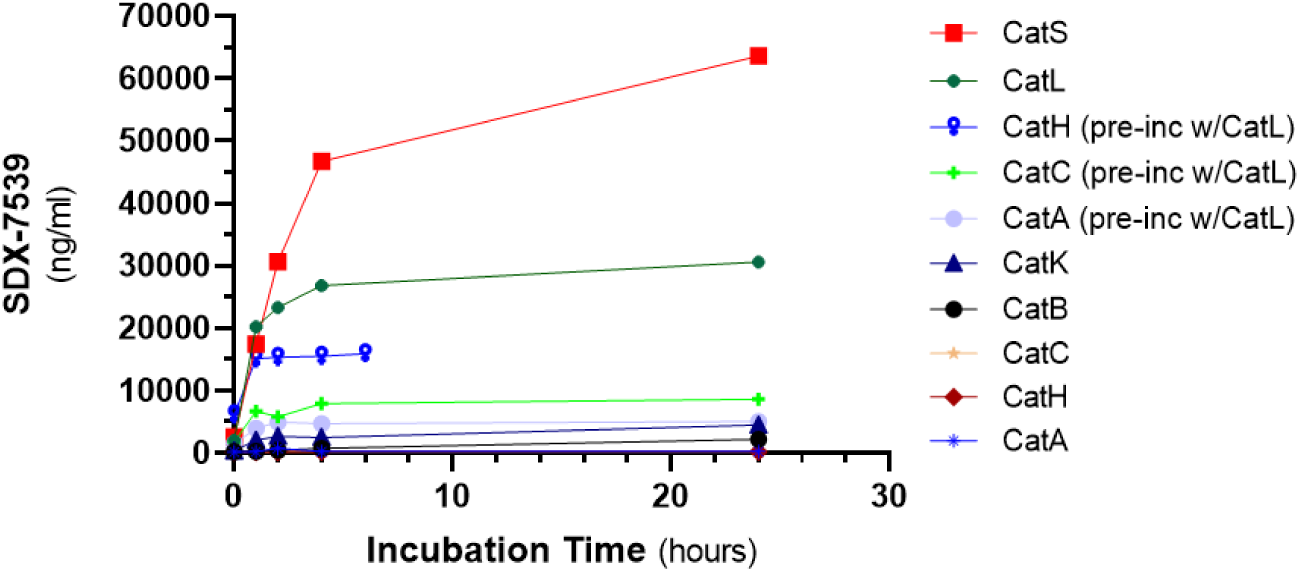
Enzyme-Mediated Metabolism of SDX-7320 *in vitro*. SDX-7320 was incubated (0.5 mg/mL) with the indicated enzymes at 37°C and SDX-7539 was measured at various time points using LC/MS. Quantitation of SDX-7539 was achieved using a deuterated internal standard (LLOQ = 48 ng/mL, ULOQ = 200,000 ng/mL).

### SDX-7539 selectively binds to MetAP2 and inhibits its catalytic activity

The binding of SDX-7320, SDX-9402 (the salt form of SDX-7539) and previously studied fumagillin-class MetAP2 inhibitors to MetAP2 was measured *in vitro* by competition ELISA (33). The binding of CKD-732 was less potent than SDX-7539 while SDX-9246 (N-acetyl analog of SDX-7539) was similar to SDX-9402 (Figure S2). Importantly, both the PDC SDX-7320 and the small molecule SDX-9178 (lacking the spiroepoxide warhead present in SDX-7539) were inactive in this assay. These results support the established role of the spiroepoxide in binding of small molecule fumagillol analogs to MetAP2 and also demonstrate that SDX-7539, when conjugated to the polymeric backbone, is restricted from binding to MetAP2.

Off-target activity of the salt form of SDX-7539 (SDX-9402) was assessed against a panel of receptors/transporters and enzymes (n=38 and 6, respectively). No effect was observed at any of the receptors or enzymes studied (up to a concentration of 10 μM SDX-7539; Tables S3 and S4).

The interaction of SDX-7539 with MetAP2 was studied further by creating co-crystals of SDX-7539 and MetAP2, followed by x-ray crystallography. The structure of the co-crystals was resolved to 1.6 Å and demonstrated covalent modification of active site histidine-231 (Figure 3, Table S5). There is a high degree of overlap between the structure of SDX-7539 bound to MetAP2 and the reported structure of human MetAP2 co-crystallized with TNP-470 (Figure S3).

**Figure 3.**
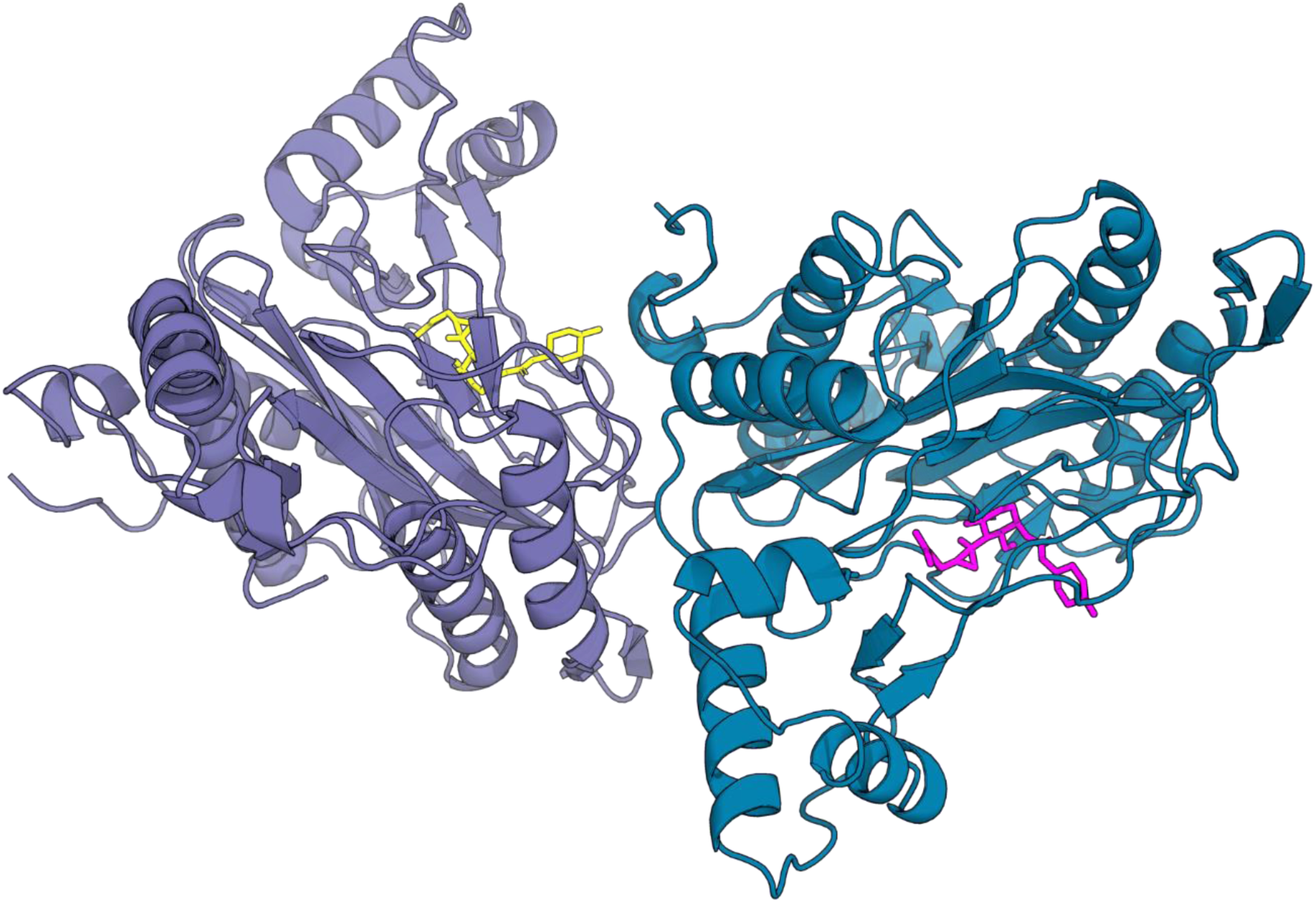
X-ray Crystal Structure of MetAP2 Complexed with SDX-7539. MetAP2 (purple and blue) forms a crystallographic dimer with both protein molecules bound to SDX-7539 (yellow and magenta).

Inhibition of aminopeptidase enzyme activity *in vitro* demonstrated both potency and selectivity of SDX-7539 for MetAP2 over MetAP1 (Figure 4, Figure S4). SDX-7539 was equipotent compared to TNP-470 against MetAP2, while neither compound inhibited MetAP1 (up to 1000 nM, Figure S4). Both of these fumagillin-class MetAP2 inhibitors exhibited time-dependent shifts in IC_50_, a hallmark of irreversible enzyme inhibitors. The PDC SDX-7320 was inactive against MetAP2 (up to 10 μg/ml) in agreement with findings from competition binding assays (Figure 4G). Note that the apparent inhibition observed at 10 μg/ml SDX-7320 was also observed in the absence of MetAP2 in the enzyme assay. The analog SDX-9178, which lacks the spiroepoxide warhead, was inactive against MetAP2, underscoring the importance of this functional group for inhibition of MetAP2 (Figure 4H).

**Figure 4.**
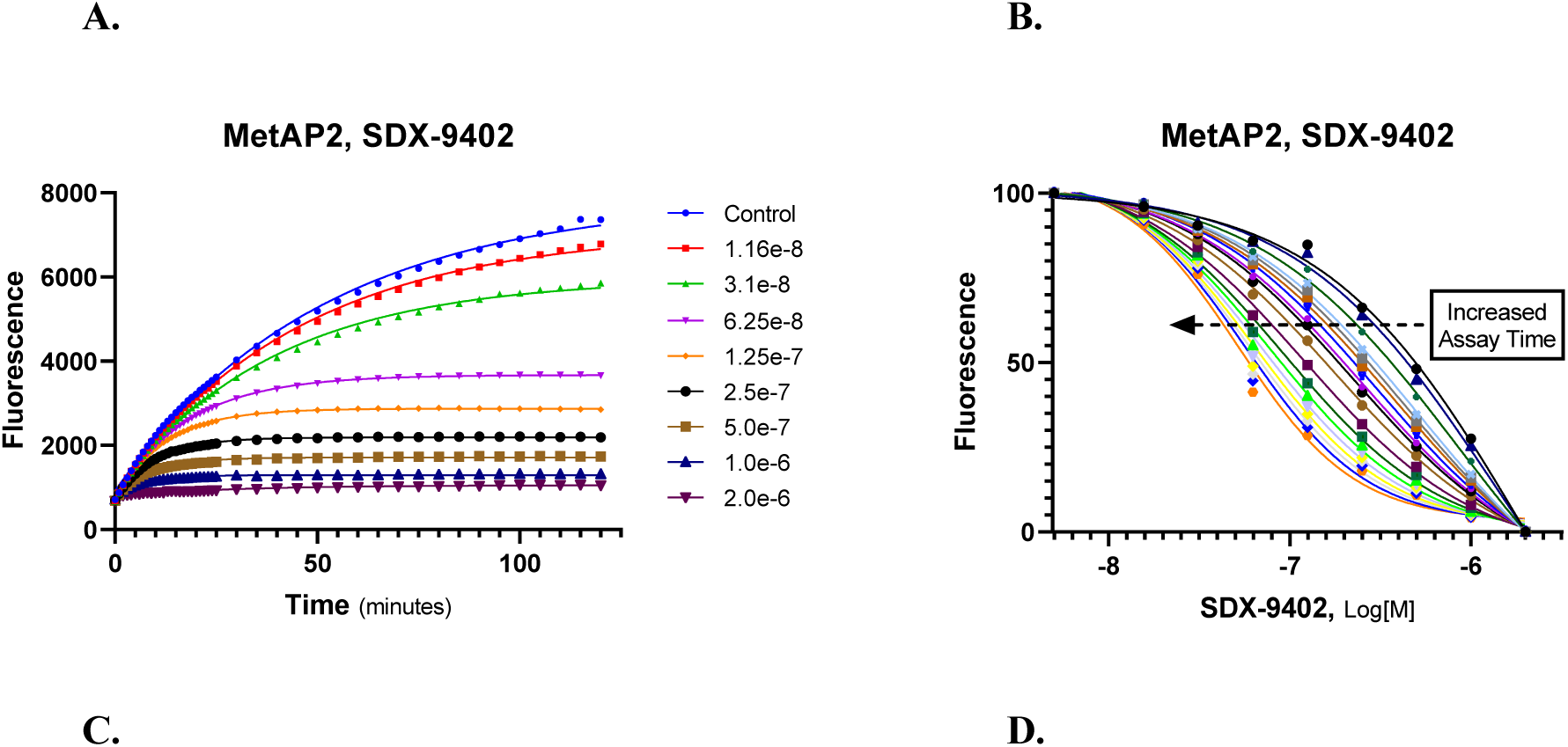
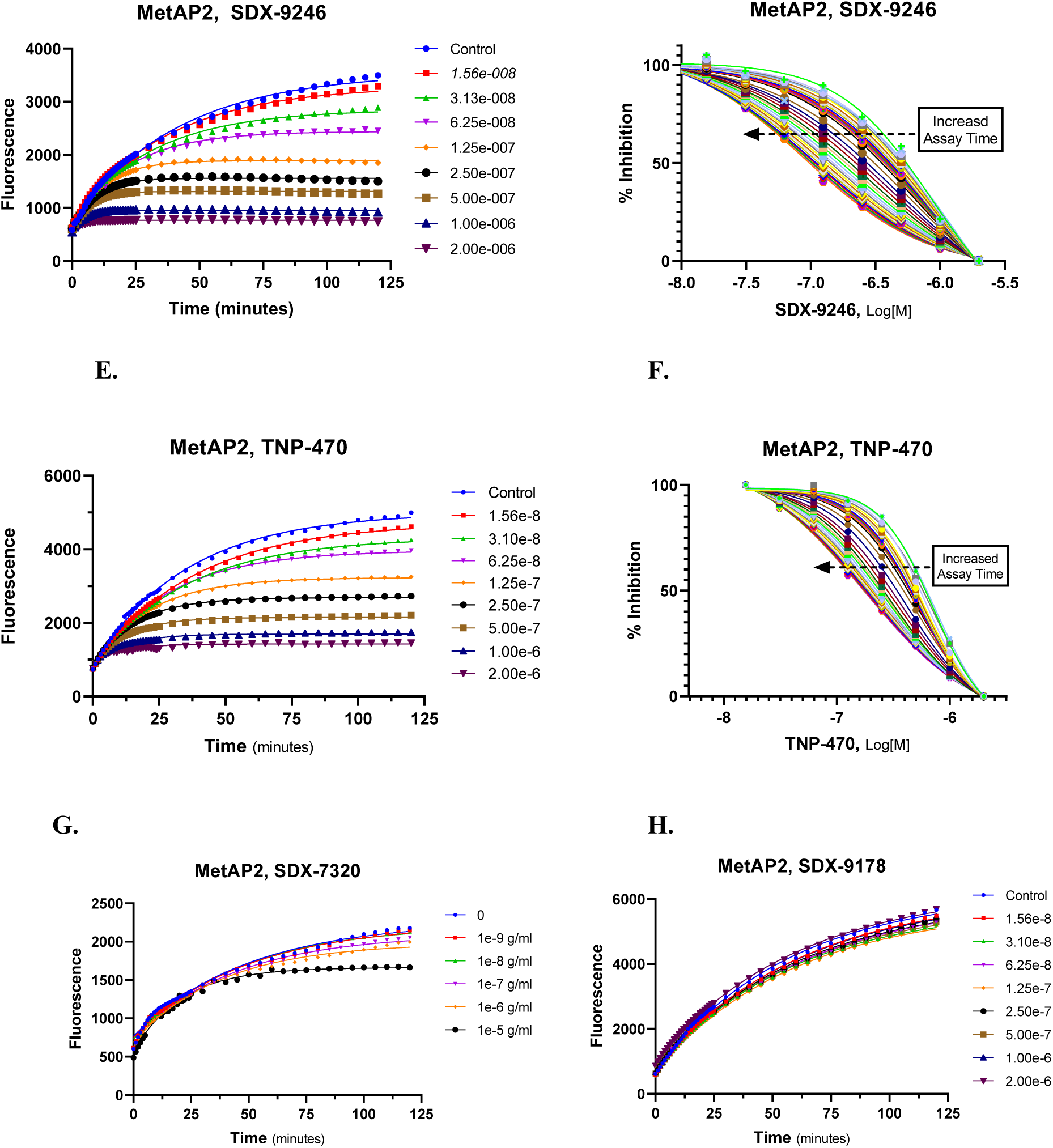
Inhibition of MetAP2 Aminopeptidase Activity. The aminopeptidase activity of human MetAP2 was measured in the presence of SDX-9402 (besylate salt of SDX-7539), SDX-9246 (N-acetyl analog of SDX-7539), TNP-470, SDX-7320 or SDX-9178 (inactive metabolite of SDX-7539). Compounds were diluted in DMSO, then added to reactions containing human recombinant MetAP2 plus the artificial fluorescent substrate Met-AMC. Fluorescence was monitored for the indicated times at room temperature and data were analyzed using GraphPad Prism v9.0. Note that the IC_50_ values for each compound decreased over time (i.e., increase in potency), characteristic of irreversible inhibitors.

### Effect of SDX-7320, SDX-7539 and TNP-470 on endothelial cell growth

Cytostatic inhibition of endothelial cell growth is a hallmark of fumagillin and its analogs (12,34). SDX-7539 and SDX-7320 both inhibited the growth of HUVECs (Figure 5). SDX-7539 inhibited growth of HUVECs with sub-nM potency while TNP-470 was approximately 3-fold less potent than SDX-7539. The PDC SDX-7320 inhibited the growth of HUVECs, albeit with >500-fold reduced potency relative to the small molecule inhibitors. As a prodrug, the decreased potency of SDX-7320 is likely due to the requirement for cellular uptake followed by lysosomal processing to release the small-molecule MetAP2 inhibitor SDX-7539. The small molecule analog SDX-9178 (which lacks MetAP2 binding and enzyme inhibition) did not affect cell growth (up to 1000 nM), indicating the requirement for MetAP2 inhibition within this chemical series to decrease the growth of HUVECs (not shown).

**Figure 5.**
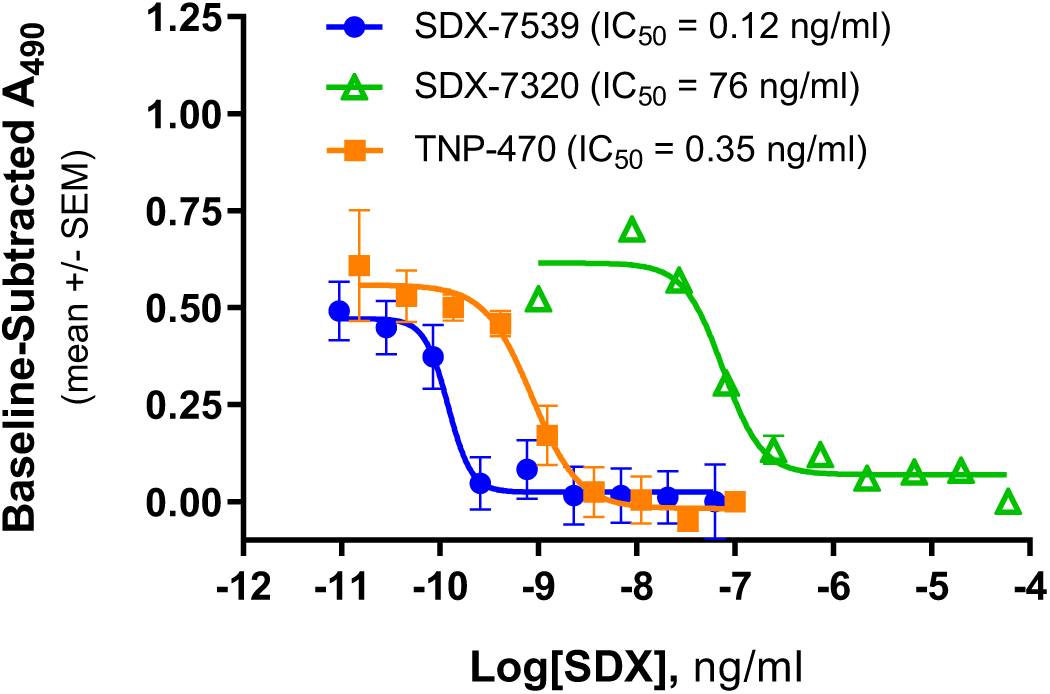
Inhibition of HUVEC Growth by SDX-7320, SDX-7539 and TNP-470. Growth of HUVECs was measured (CellTiter 96^®^, Promega) after 72 hours incubation in the presence and absence of the indicated compounds. IC_50_ values were calculated by non-linear regression (GraphPad Prism v9.0) after subtracting absorbance measured in the absence of test compounds.

### CNS safety and pharmacokinetic characterization of SDX-7320 and related compounds in rats

The pharmacokinetic profile of the small molecule SDX-7539 was assessed in rats, following IV bolus as the unconjugated small molecule and as the PDC SDX-7320 (Figure S5). The terminal elimination half-life for SDX-7539 after IV administration ranged from 10-15 minutes as the unconjugated small molecule, whereas after dosing the PDC SDX-7320, the half-life of SDX-7539 was calculated to be approximately 10 hours, most likely due to the additional processing required for enzymatic release of SDX-7539 from the PDC.

A primary driver for conjugating the small molecule MetAP2 inhibitor SDX-7539 to a high molecular-weight polymer was to limit CNS exposure and minimize CNS-related toxicity. To assess the impact of polymer conjugation on CNS exposure, levels of small molecules (SDX-7539, TNP-470) in plasma and cerebrospinal fluid (CSF) were measured after IV dosing of either SDX-7320, SDX-7539 or TNP-470 at equimolar small molecule doses in rats. Since TNP-470 undergoes rapid metabolism *in vivo*, producing an active metabolite (M-IV), the M-IV metabolite was measured as opposed to the parent molecule. CSF levels of SDX-7539 (measured near plasma T_max_) following IV administration of SDX-7320 were >50-fold lower compared to when an equivalent amount of free (unconjugated) SDX-7539 was administered, and 17-fold lower than the active metabolite of TNP-470 (Table S6).

To assess the CNS side effect profile of SDX-7320 *in vivo*, the neurobehavioral FOB was conducted in rats under GLP conditions, at 12 and 36 hours after a single SC dose of SDX-7320 (3, 10 and 30 mg/kg). No signs of abnormal CNS-related behavioral toxicity (e.g., tonic movements, startle-response, hind-limb splay) were observed in rats up to 36 hours following treatment with SDX-7320 (Table S7). A decrease in body weight gain relative to vehicle-treated animals was found following dosing with 10, and 30 mg/kg SDX-7320 (36 hours post-dose). This finding was not considered adverse due to the slight magnitude of the changes (<5%), and is not unexpected given the known effects of this drug class on adipose tissue and body weight (12, 36). There were no other SDX-7320-related changes during the study.

### Effect of SDX-7320 on tumor growth and metastasis

Anti-tumor activity of SDX-7320 was initially evaluated in A549 xenografts, a model of NSCLC and was compared to the small molecule MetAP2 inhibitors SDX-7539 and TNP-470, dosed at equimolar small molecule doses. SDX-7320, administered IV (6, 60 mg/kg, SC, Q4D), caused a significant, dose-dependent inhibition of A549 tumor growth, relative to vehicle-treated controls (Figure 6A). Tumor growth inhibition in response to 6 mg/kg SDX-7320 (IV, every 4D, which delivers the equivalent of 1.2 mg/kg SDX-7539 at each dose) matched what was observed with the small molecule SDX-7539 (IV, 37 mg/kg, every 2D) and exceeded the efficacy observed in response to TNP-470 (IV, 30 mg/kg, every 2D), highlighting the potential advantage of polymer conjugation from a dose-effect and dosing schedule perspective.

**Figure 6.**
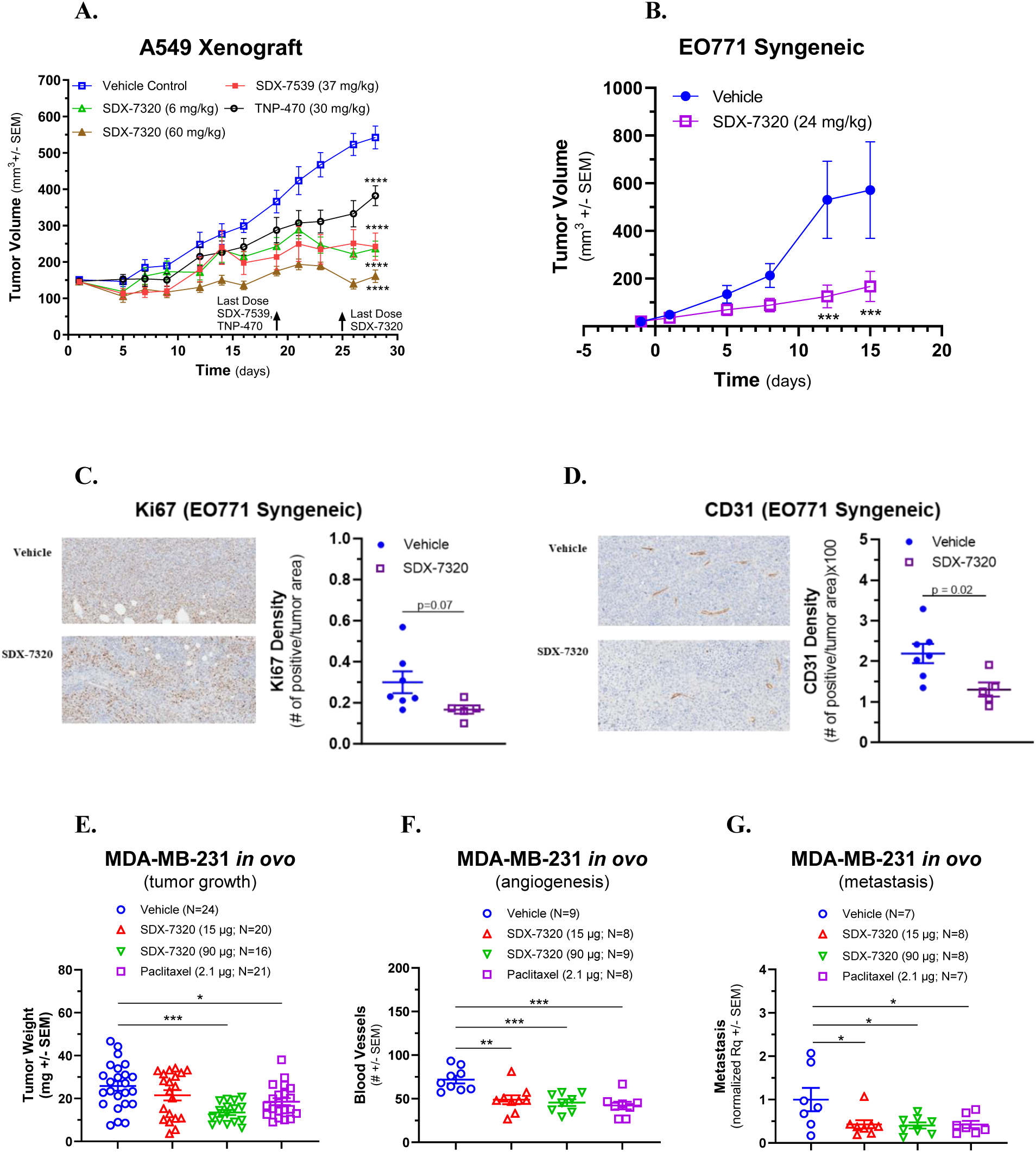
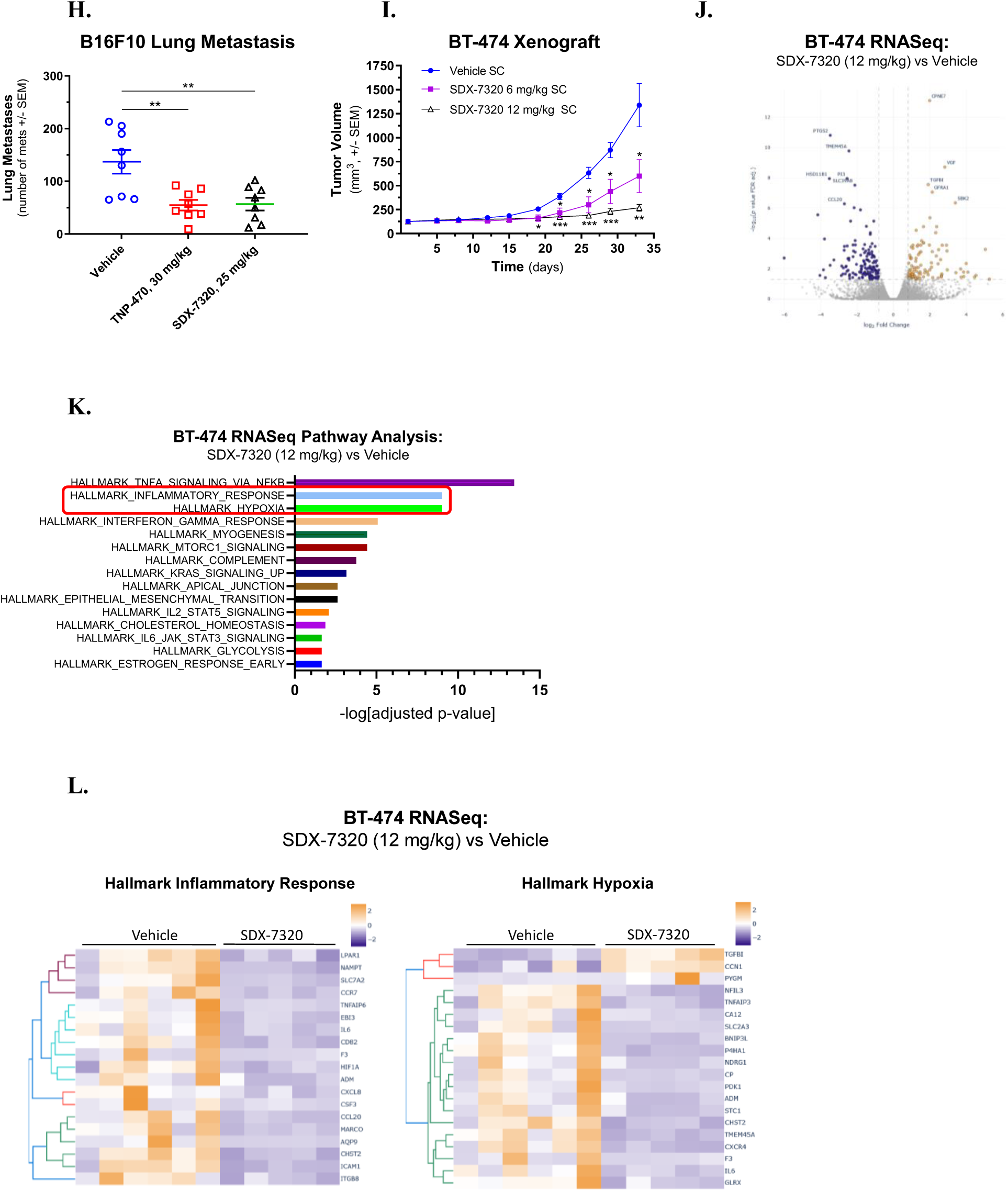
Anti-Tumor Activity of SDX-7320 and SDX-7539 in Xenograft and Syngeneic Models of Cancer. **A.** A549 xenograft model: SDX-7320 and SDX-7539 were injected IV, Q4D or QOD, respectively, into nude mice (ten/group) with established A549 tumors (treatment was initiated when tumors were between 100 and 200 mm^3^). TNP-470 was injected IV every other day. Statistical analysis was carried out by two-way ANOVA with Sidack’s multiple comparisons test (GraphPad,v9.0). **B.** EO771 syngeneic model of TNBC: ovariectomized mice with orthotopic EO771 tumors were dosed with SDX-7320 (Eight mice/group, 24 mg/kg, SC, Q4D; total of four doses) after which tumors were dissected and fixed in buffered formalin for histology and IHC. Statistical analysis of tumor volume was carried out by two-way ANOVA with Sidack’s multiple comparisons test; N=8 mice/group (GraphPad,v9.0). **C, D.** Immunostaining of formalin-fixed, paraffin-embedded tumor tissue with antibodies against Ki67 or CD31. The density of staining (# of positive cells/tissue area) was quantified using Aperio software (Leica Inc). Statistical analysis was carried out by unpaired t-test (GraphPad,v9.0). **E.** MDA-MB-231 tumors were grown in chick embryos (inoculated on day E9, then dissected and weighed on day E19). Treatment with vehicle, paclitaxel (2.1 μg, every other day, four doses), SDX-7320 (15, 90 μg, every four days, two doses) was initiated on day E11. Statistical analysis was conducted using one-way ANOVA with Dunnett’s multiple comparisons test (GraphPad v9.0) **F.** MDA-MB-231 tumor blood vessel density on day E18 for vehicle, paclitaxel (2.1 μg), SDX-7320 (15, 90 μg). **G.** MDA-MB-231 tumor metastasis into the lower CAM after treatment with vehicle, paclitaxel (2.1 μg, QOD, four doses), SDX-7320 (15, 90 μg, every four days, two doses). **H.** B16-F10 syngeneic model of lung metastases: treatment with vehicle, TNP-470 (30 mg/kg, IP, QOD) or SDX-7320 (25 mg/kg, SC, Q4D) was initiated one day after tumor cells were administered *via* tail vein injection (eight mice/group). **I.** BT-474 xenograft tumor growth (model of HR+/HER2+ breast cancer; eight mice/group) after SDX-7320 (6, 12 mg/kg, SC, every four days). Statistical analysis of tumor volume was conducted using a mixed-effects model with Dunnett’s multiple comparisons (GraphPad v9.0) **J.** Volcano plot of differentially expressed genes (DEGs) from tumors in SDX-7320 (12 mg/kg)-treated mice (N=5) relative to tumors from vehicle-treated mice (N=6). **K.** Results of pathway enrichment analysis (Enricher) conducted on DEGs (n=268) from SDX-7320 (12 mg/kg)-treated tumors relative to tumors from vehicle-treated mice (only pathways with enrichment scores with p≤0.01 are shown). **L.** Heat maps of the top 20 DEGs from two selected pathways (Hallmark Inflammatory Response, Hallmark Hypoxia) that were significantly altered in tumors from SDX-7320-treated mice.

To further investigate the potential anti-tumor effects of SDX-7320, EO771 allografts, a model of TNBC were used. SDX-7320 (24 mg/kg SC, Q4D) significantly inhibited tumor growth (Figure 6B), and histological analysis of dissected tumors showed reduced staining for Ki67 and CD31, indicating anti-proliferative and anti-angiogenic activity, respectively (Figure 6C and D).

SDX-7320 was then tested in a model of human TNBC using MDA-MB-231 tumors grown in chick embryos with paclitaxel serving as a positive control. SDX-7320 (15 and 90 μg, every 4D) was administered twice, beginning two days after implantation of tumor cells. The results demonstrated dose-dependent inhibition of tumor growth relative to vehicle-treated tumors, with the higher dose exceeding the efficacy of paclitaxel (2.1 μg, every other day, total of 4 doses, Figure 6E). The effect of SDX-7320 on angiogenesis was assessed two days prior to termination, and a significant decrease in blood vessel density was observed (Figure 6F).

Because the chick embryo tumor model has been used to determine whether a drug possesses anti-metastatic activity (35), the anti-metastatic effects of SDX-7320 were determined in this model. Quantitation of human DNA in the lower chorioallantoic membrane was used as a measure of metastasis, and the results showed that SDX-7320 significantly inhibited metastasis of MDA-MB-231 tumors (Figure 6G).

Building upon the findings in the chick embryo model of metastasis, the activity of both SDX-7320 and TNP-470 were investigated in a syngeneic B16-F10 melanoma lung metastasis assay. Drugs were administered beginning one day after tumor cells were introduced *via* tail vein injection. SDX-7320 significantly inhibited the outgrowth of B16-F10 melanoma cells in the lungs relative to vehicle-treated animals, and an identical degree of inhibition was observed with TNP-470 albeit at a dose that was six times greater than the dose of SDX-7320 (comparison is based upon small-molecule equivalents; Figure 6H).

To gain additional mechanistic insight into the anti-tumor effects of SDX-7320, BT-474 cells were utilized in an orthotopic xenograft model of HR+/HER-2+ breast cancer. SDX-7320 elicited significant, dose-dependent inhibition of BT-474 tumor growth (6, 12 mg/kg; Figure 6I). Next RNA-seq was used to compare gene expression in tumors from vehicle-treated versus SDX-7320-treated mice (12 mg/kg). Principal component analysis (PCA) of the RNASeq results showed that gene expression in vehicle samples and in SDX-7320-treated samples each clustered with their own groups, however expression differences across principal component one indicated one vehicle sample may have significantly different gene expression than the other samples within this group (Figure S6). A number of genes were identified with significant differences in expression between vehicle-treated and SDX-7320-treated tumors (Figure 6J). Pathway analysis conducted on the set of genes with significant differences in expression between the two groups showed that several pathways important in tumor biology including “Hallmark Inflammatory Response” as well as “Hallmark Hypoxia” were significantly altered in tumors from SDX-7320 vs. vehicle-treated mice (Figure 6K). The expression of several hypoxia-regulated genes including the master regulator of the hypoxic response, HIF1α, were significantly reduced in tumors from SDX-7320-treated mice (Figure 6L). This suggests that inhibiting the adaptive response of BT-474 tumors to hypoxia is one mechanism by which SDX-7320 inhibits tumor growth, or alternatively that normalization of tumor vasculature following inhibition of MetAP2 led to the attenuation of hypoxia (36). Genes related to inflammation were significantly affected in tumors from SDX-7320-treated mice, including interleukin-6, chemokine genes CXCL1, CXCL3, CXCL8 and CCL20, as well as the chemokine receptors CCR7 and CXCR4 which were all significantly suppressed (Figure 6L and Supplementary Data DEG file).

## Discussion

The ability of the mycotoxin fumagillin, and semi-synthetic derivatives such as TNP-470, to act as irreversible inhibitors of MetAP2, with consequent anti-angiogenic and anti-proliferative potential, is well established (37). TNP-470 itself has been extensively investigated and has displayed potent anti-tumor activity in multiple preclinical models as well as in the clinical setting (24,25,38). A poor PK profile and limited tolerability of TNP-470 have driven the exploration of structural modifications based on the fumagillol pharmacophore, retaining the irreversible nature of MetAP2 inhibition (1). SDX-7539 is a novel small molecule fumagillol derivative bearing a 4-aminocyclohexyl carbamate with *trans* stereochemistry across the cyclohexane ring whose half-life is 4-fold longer than that of TNP-470, and does not possess the *N*-oxidation potential nor the hydrolytic ester lability *in vivo* of some second-generation analogs (e.g., CKD-732) (39,40) (Figure 1). SDX-7539 is a potent and selective inhibitor of MetAP2, based on results from *in vitro* binding, enzyme and cell-based assays. X-ray structure determination of MetAP2-SDX-7539 co-crystals yielded strong similarities to the published structures of MetAP2-Fumagillin and MetAP2-TNP-470, confirming the nucleophilic attack of the histidine-231 imidazole on the spiroepoxide, forming a covalent bond in the enzyme active site (41).

Polymer-conjugation has been investigated for several decades, mainly with the aim of enhancing drug delivery and improving the therapeutic index of pharmacologically active small molecules (26). Covalent conjugation to various water soluble polymers using TNP-470 as starting material has been reported wherein facile nucleophilic displacement of the TNP-470 chloride heteroatom was utilized to generate anti-angiogenic PDCs bearing a 2-*O*-acetyl-linked carbamoylfumagillol attached to poly(vinyl alcohol), or bearing 2-aminoethylglycyl-carbamoylfumagillol attached to poly(HPMA) *via* a tetrapeptide linker (29,42). Although these PDCs showed promising preclinical activity in their respective indications, neither proceeded to human clinical evaluation. The PDC SDX-7320 is based on a non-immunogenic, biocompatible, hydrophilic p(HPMA) polymer backbone (43,44) and utilizes a glycyl-L-phenylalanyl-L-leucyl-glycyl-(GFLG) tetrapeptide linker to conjugate the selective SDX-7539 MetAP2 inhibitor (Figure 1).

In the genesis of the high molecular weight polymer conjugate SDX-7320 and its active component SDX-7539, objectives were: to ameliorate the dose-limiting CNS toxicity of historical fumagillin derivatives such as TNP-470 by reducing the amount of small molecule penetrating the blood brain barrier; to provide a sustained PK profile of the active moiety suitable for convenient administration; and to harness potential preferential tumor accumulation of a novel, potent MetAP2 inhibitor *via* the enhanced permeability and retention effect (EPR) effect. This report describes the characteristics of SDX-7320 and its pharmacological profile as a novel anti-tumor agent.

In terms of physical properties, as described above the active pharmaceutical ingredient SDX-7320 has high aqueous solubility (>200 mg/mL) and excellent long-term chemical stability (> 48 months) has been observed at ambient conditions (stability data not shown). On a weight basis, the SDX-7539 payload was determined to be approximately 20%, and the SDX-7320 average molecular weight (M_w_) was approximately 30 kDa, with a particle size distribution measured to be 10 nm Z-average (PDI < 0.25) in aqueous solution. Accordingly, SDX-7320 is a high drug loading, hydrophilic PDC which has the potential to take advantage of preferential macromolecular passive tumor accumulation while remaining largely within the approximate glomerular filtration threshold (about 30-50 kDa) suitable for renal clearance (45). Importantly, free, unconjugated SDX-7539 was not detected in SDX-7320 as isolated, and based on the relative activities of the PDC versus the small molecule, MetAP2 inhibition is not achieved until SDX-7539 is released enzymatically.

The tetrapeptide linker “GFLG” has been previously used to construct PDCs and can be hydrolyzed by lysosomal proteases such as cathepsin B (CTSB, (46)). However, as endopeptidase activity is required, the proximal structure of the drug being cleaved is clearly fundamental to isoform specificity and efficiency: only select cysteine cathepsin isoforms were capable of releasing SDX-7539 from SDX-7320, with Cathepsin S (CTSS) exhibiting the highest activity *in vitro* against SDX-7320.

In the case of SDX-7320 a major objective of polymer conjugation was to improve the therapeutic index relative to previous fumagillin-based MetAP2 inhibitors such as TNP-470 by conjugating the pharmacologically active moiety SDX-7539 to a high molecular weight prodrug backbone, thereby limiting CNS exposure. Analysis following IV administration of TNP-470, SDX-7539 or the PDC SDX-7320 at equimolar small molecule doses demonstrated a marked reduction in CSF levels of SDX-7539 when dosed as the PDC compared to the unconjugated small molecule or to TNP-470 (Table S6). Additionally, no significant behavioral abnormalities were observed in rats following a single doses of the SDX-7320 PDC in a GLP functional observational battery (Table S7).

As a macromolecular system structurally based on poly(HPMA) with a M_w_ ∼ 30 kDa, SDX-7320 is purported to benefit from two passive tumor-, and tumor microenvironment-targeting processes: the EPR effect and extravasation through leaky vasculature and inflammatory cell-mediated sequestration (ELVIS). The EPR effect is a well-described, albeit mechanistically controversial, concept which may enable appropriately sized nanoparticles to preferentially accumulate, relative to small molecules, in solid tumors *via* diffusion through the complex, immature neovasculature with reduced escape due to limited lymphatic drainage (47,48). ELVIS is a more recently described phenomenon in which certain HPMA-based copolymer systems have been reported to target tumor-associated macrophages (49,50).

Polymer conjugation is anticipated to be most beneficial when conjugating inhibitors with high potency against the target of interest, in order to limit the total dose of PDC. Unexpected reductions in efficacious doses of the SDX-7320 PDC were observed compared to either the released small molecule SDX-7539 alone or other small molecule MetAP2 inhibitors, in xenograft and in syngeneic models of cancer (Figure 6A and H). For example, in the B16-F10 melanoma model (Figure 6H) when SDX-7320 was dosed at 25 mg/kg (delivering approximately 5 mg/kg SDX-7539 per dose) every four days, it was equally efficacious as TNP-470 dosed at 30 mg/kg every other day. After 15 days of treatment, the effect on lung metastases in response to each agent was virtually identical despite an 83% reduction in total dose of small molecule MetAP2 inhibitor delivered with the PDC. Polymer-conjugation likely contributes to the reduced total drug dose required for efficacy by prolonging MetAP2 inhibition, secondary to increased half-life of the inhibitor, and possibly by enhancing tumor uptake, due to the EPR and ELVIS effects. To the best of our knowledge, this is the first instance that reductions in small molecule-equivalent efficacious doses following polymer conjugation have been reported, as typically the total dose of small molecule equivalent increases following polymer conjugation, rendering SDX-7320 unique in this respect.

The mechanism by which SDX-7320 inhibits tumor growth is multi-factorial, extending beyond its anti-angiogenic effects, to include inhibition of metastases. Analysis of gene expression data from BT-474 tumors treated with SDX-7320 identified multiple pathways that were affected, including hypoxia and inflammatory responses which may contribute to inhibition of tumor growth following MetAP2 inhibition.

This report represents the initial disclosure of the structure, synthesis and preclinical pharmacological profile of the polymer drug conjugate SDX-7320, and its active payload SDX-7539. As a polymer-based prodrug, SDX-7320 has displayed multiple improved characteristics compared to historical fumagillin-derived small molecule analogs such as TNP-470. This MetAP2 inhibitor, a next generation nanomedicine, has completed a Phase 1 clinical dose-escalation safety trial in solid tumors (NCT02743637) and is currently being investigated in multiple Phase 1b/2 clinical proof-of-concept studies in patients with triple-negative breast cancer and hormone receptor-positive breast cancer (NCT05570253, NCT05455619).

## Acknowledgements

The authors would like to acknowledge the seminal work of Dr Judah Folkman in the field of angiogenesis, Dr Donald Ingber, MD, PhD in discovering the anti-angiogenic activity of fumagillin, Dr Daniel Von Hoff for discussions in the development of SDX-7230 and Ronit Satchi-Fainaro, PhD for her early work on polymer conjugates. The following are acknowledged for executing preclinical experiments and data collection: SBH Sciences (Natick, MA), Neosome Life Sciences (Woburn, MA, Sara Little), QPS (Newark, DE), Vivisource (Lexington, MA), Apredica (Watertown, MA), MPI Research (Mattawan, MI), TD2 (Scottsdale, AZ), Inovotion (Grenoble, France), Wax-It Histology Services (Vancouver, BC, Canada), Azenta/GeneWiz (Waltham, MA), Watershed AI (Cambridge, MA), Pharm-analyt GmbH (Baden, Austria), Peak Proteins (Macclesfield, UK, Dr Stephen Moss). Additionally, the following are acknowledged for performing chemical synthesis and analytical development: Organix, Inc. (Woburn, MA, Dr Paul Blundell, Dr Mario Gonzales, Dr Jie Li), Solid Form Solutions Ltd. (Milton Bridge, Scotland), PCI, Inc. (Newburyport, MA), Lyophilization Services of New England, Inc. (Bedford, NH), Omm Scientific, Inc. (Dallas, TX, Dr Donald Stewart), Boston Analytical (Salem, NH), Jordi Labs, LLC. (Bellingham, MA), HepatoChem, Inc. (Beverly, MA), Patheon (Ferentino, Italy), Custom NMR Services (Woburn, MA, Dr Jin Hong) and CuriRx (Wilmington, MA). The authors would like to thank Andrea Gwosdow and Becca Kranz (Gwosdow Associates Science Consultants) for their assistance in preparation of the draft and are grateful to Prof. Bruce Zetter, PhD, Dr Andrew J. Dannenberg and Dr Rakesh Jain for providing feedback on the manuscript.

