## Supplemental Data for "Pharmacological Characterization of SDX-7320/Evexomostat: a Novel Methionine Aminopeptidase Type 2 Inhibitor with Anti-Tumor and Anti-Metastatic Activity"

**Supplementary Data: Materials, Methods and Results**

**Table of Contents**

**Supplemental Materials and Methods**

Chemistry methods ………………………………………………………………………………...3

Enzyme-mediated metabolism of SDX-7320 *in vitro* ……………………………………………...9

Expression of MetAP2 and MetAP2-SDX-7539 co-crystal structure …………………………......9

Growth of MDA-MB-231 tumors in chick embryos ……………………………………………..12

**Supplemental Figures**

Figure S1. SDX-7320 atom numbering for NMR assignments……………………………………..5

Figure S2. *In vitro* competition binding to MetAP2 ……………………………………………...14

Figure S3. Superposition of crystal structures MetAP2-SDX-7539 and MetAP2-TNP-470 …….14

Figure S4. MetAP1 enzyme assay ………………………………………………………………..15

Figure S5. Pharmacokinetics of SDX-7539 in rats …………………………………………….....15

Figure S6. Principle component analysis (PCA) of BT-474 RNASeq Data ……………………..16

**Supplemental Tables**

Table S1. ^1^H and ^13^C Resonance assignments …………………………………………………….6

Table S2. Commercial sources of cathepsins ……………………………………………………..17

Table S3. Measurement of off-target binding of SDX-9402 ……………………………………..17

Table S4. Measurement of off-target enzyme inhibition by SDX-9402 ...……………………….20

Table S5. Data collection, refinement statistics - MetAP2-SDX-7539 co-crystal structure ……..20

Table S6. CNS Exposure following peripheral dosing of small molecules and SDX-7320 ...…...21

Table S7. Results of the rat FOB ………………………………………………………………....22

**Supplementary References** ……………………………………………………………………..23

**Chemistry methods**

Fumagill-6-yl *p*-nitrophenyl carbonate (CAS# 1160640-95-8) was prepared from commercially available fumagillin dicyclohexylamine (CAS# 41567-78-6) based on published methods (1). The synthesis of glycine, *N*-(2-methyl-1-oxo-2-propen-1-yl)glycyl-L-phenylalanyl-L-leucyl-, 4-nitrophenyl ester, (CAS# 100424-71-3) and *N*-(2-hydroxypropyl)methacrylamide, HPMA, (CAS# 21442-01-3) is described in literature procedures (2,3). TNP-470 and CKD-732 were purchased from commercial suppliers or prepared based on published procedures (4,5). Synthesis of SDX-9402 (salt form of free base SDX-7539, CAS# 1631953-52-0), SDX-7320 (CAS# 2416263-67-5), SDX-9246, SDX-9280 and SDX-9178 are described below.

**Fumagill-6-yl *N*-(*trans*-4-aminocyclohexyl)carbamate benzenesulfonic acid salt [(1*R*,4*r*)-4-(((((3*R*,4*S*,5*S*,6*R*)-5-Methoxy-4-((2*R*,3*R*)-2-methyl-3-(3-methylbut-2-en-1-yl)oxiran-2-yl)-1-oxaspiro[2.5]octan-6-yl)oxy)carbonyl)amino)cyclohexan-1-aminium benzenesulfonate] (SDX-9402):**

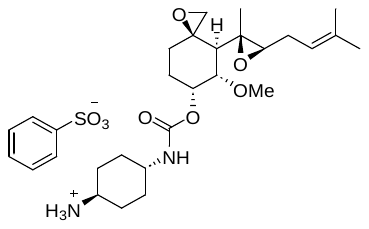

*trans*-1,4-Diaminocyclohexane (102.2 g, 0.895 mol) was dissolved in ethyl acetate (2.8 L) and cooled to 0-5^o^C. Fumagill-6-yl *p*-nitrophenyl carbonate (CAS# 1160640-95-8; 80.8 g, 0.181 mol) was dissolved in ethyl acetate (3.2 L), added to the reaction mixture and stirred for 2 h at 0-5^o^C. The mixture was filtered and the solids rinsed with ethyl acetate (2 x 1.2 L). The filtrate was washed with water (0.5 L) and extracted with a mixture of water (1.6 L) and 10% citric acid (0.24 L). 1N Aqueous sodium carbonate (0.87 L) was added to the aqueous layer and the free base was extracted into MTBE. The organic phase was dried (sodium sulfate), filtered and washed with MTBE. The solution was cooled to 0-5^o^C and seeded with fumagill-6-yl *N*-(*trans*-4-aminocyclohexyl)carbamate benzenesulfonic acid salt (0.8 g). A solution of benzenesulfonic acid (17.7 g, 0.112 mol) in MTBE (0.43 L) was added and the mixture was cooled to -20^o^C. The solids were filtered, washed with MTBE (0.5 L) and dried under vacuum to yield fumagill-6-yl *N*-(*trans*-4-aminocyclohexyl)carbamate benzenesulfonic acid salt (68.3 g, HPLC 99.0% w/w). Methanol (200 mL) was added and the mixture was heated to 40^o^C. MTBE (250 mL) was charged and the mixture was cooled to 35^o^C. Seeds of fumagill-6-yl *N*-(*trans*-4-aminocyclohexyl)carbamate benzenesulfonic acid salt were charged and the mixture was cooled to 2-8^o^C. MTBE (330 mL) was added and the mixture was stirred at 2-8 ^o^C for 15 h. The solids were collected by filtration, washed with MTBE, and dried under vacuum to provide fumagill-6-yl *N*-(*trans*-4-aminocyclohexyl)carbamate benzenesulfonic acid salt (64.6 g, 62% yield) as a white crystalline powder: HPLC (Method 1) 99.9% w/w; *m/z* (APCI^+^) 423.2 [M + H]^+^ 100%; v_max_ (cm^-1^) 3324, 2949 (br), 1711, 1536; ^1^H NMR (400 MHz, CDCl_3_) δ 1.01-1.10 (3H, br-m), 1.19 (3H, s), 1.39-1.47 (2H, br-m), 1.65 (3H, s), 1.74 (3H, s), 1.78-1.82 (1H, m), 1.90-2.07 (7H, m), 2.10-2.18 (1H, m), 2.32-2.37 (1H, m), 2.52 (1H, d, *J* 6.2 Hz), 2.56 (1H, a-t, *J* 6.2 Hz), 2.89-2.94 (2H, m), 3.22-3.36 (1H, m), 3.43 (3H, s), 3.62 (1H, dd, *J* 11.2 Hz, *J* 2.8 Hz), 4.99 (1H, br-s), 5.18 (1H, a-t, *J* 7.2 Hz), 5.42 (1H, br-s), 7.40-7.44 (3H, m), 7.70 (3H, br-s), 7.85 (2H, dd, *J* 5.6 Hz, *J* 2.0 Hz); ^13^C NMR (100 MHz, CDCl_3_) δ 13.74, 18.03, 25.75 (2C), 25.81, 27.34, 29.19 (2C), 30.66, 30.87, 47.99, 48.59, 49.81, 50.74, 56.58, 58.87, 59.53, 61.07, 66.69, 79.29, 118.49, 125.84 (2C), 128.59 (2C), 130.65, 134.92, 144.20, 155.25.

HPLC Method 1: HPLC spectra were obtained on an Agilent 1100 series instrument or equivalent with detection by CAD and an Agilent Zorbax Eclipse XBD C_18_ reverse-phase column (150 mm x 4.6 mm, 5 μm) at 25^o^C; flow rate 1.0 mL/min; mobile phase A, 0.1% trifluoroacetic acid in water; B, 0.1% trifluoroacetic acid in acetonitrile; gradient 5 to 95% B over 15 min; 10 μL injection volume.

**Glycine, *N*-(2-methyl-1-oxo-2-propen-1-yl)glycyl-L-phenylalanyl-L-leucyl-, 4-nitrophenyl ester, polymer with *N*-(2-hydroxypropyl)-2-methyl-2-propenamide, reaction products with (3*R*,4*S*,5*S*,6*R*)-5-methoxy-4-[(2*R*,3*R*)-2-methyl-3-(3-methyl-2-buten-1-yl)-2-oxiranyl]-1-oxaspiro[2.5]oct-6-yl *N*-(*trans*-4-aminocyclohexyl)carbamate (SDX-7320, CAS# 2416263-67-5):**

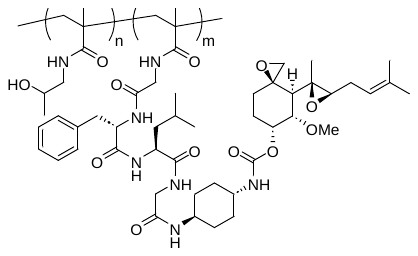

Glycine, *N*-(2-methyl-1-oxo-2-propen-1-yl)glycyl-L-phenylalanyl-L-leucyl-, 4-nitrophenyl ester (143 g, 0.246 mol) and *N*-(2-hydroxypropyl)methacrylamide, HPMA, (317 g, 2.21 mol) were stirred in acetone (4.3 L) and degassed. 2,2'-Azobis(2-methylpropionitrile) (22.4 g, 0.136 mol) was added and the mixture was heated at 50^o^C for 48 h. The mixture was cooled and acetone (1 L) was charged followed by 4-methoxyphenol (0.3 g, 0.0024 mol). The solid was isolated by filtration, washed with acetone, triturated with acetone, filtered and dried under vacuum to afford glycine, *N*-(2-methyl-1-oxo-2-propen-1-yl)glycyl-L-phenylalanyl-L-leucyl-, 4-nitrophenyl ester, polymer with *N*-(2-hydroxypropyl)-2-methyl-2-propenamide) as a white solid (365.3 g). Glycine, *N*-(2-methyl-1-oxo-2-propen-1-yl)glycyl-L-phenylalanyl-L-leucyl-, 4-nitrophenyl ester, polymer with *N*-(2-hydroxypropyl)-2-methyl-2-propenamide) (108.4 g) was dissolved in *N*,*N*-dimethylformamide (0.7 L). *N*,*N*-Diisopropylethylamine (27.3 g, 0.211 mol) was added and cooled to 5-10^o^C. Fumagill-6-yl *N*-(*trans*-4-aminocyclohexyl)carbamate benzenesulfonic acid salt (34.8 g, 0.06 mmol) was added and the mixture was warmed to room temperature and stirred for 3 h. The reaction mixture was diluted with ethyl acetate and brine. The aqueous layers were separated, concentrated under vacuum and filtered through Celite. The filter cake was washed with water and the filtrate was purified by tangential flow filtration to provide a retentate conductivity of less than 0.5 mS/cm. The aqueous solution was filtered (0.22 μm), lyophilized and the resulting solid was sieved (600 μm) to obtain glycine, *N*-(2-methyl-1-oxo-2-propen-1-yl)glycyl-L-phenylalanyl-L-leucyl-, 4-nitrophenyl ester, polymer with *N*-(2-hydroxypropyl)-2-methyl-2-propenamide, reaction products with (3*R*,4*S*,5*S*,6*R*)-5-methoxy-4-[(2*R*,3*R*)-2-methyl-3-(3-methyl-2-buten-1-yl)-2-oxiranyl]-1-oxaspiro[2.5]oct-6-yl *N*-(*trans*-4-aminocyclohexyl)carbamate (SDX-7320; 100.2 g) as an amorphous powder, as confirmed by XRPD. Potential residual unconjugated fumagill-6-yl *N*-(*trans*-4-aminocyclohexyl)carbamate (SDX-7539), acrylamide monomers, and individual unspecified unknown impurities were not detected by HPLC.

HPLC Method 2: Waters 2695 series HPLC instrument or equivalent with detection by UV; Agilent Zorbax Eclipse XBD C_8_ reverse-phase column (50 mm x 4.6 mm, 5 µm) at 30^o^C; flow rate 2.0 mL/min; mobile phase A, 0.05% phosphoric acid in water; B, 0.05% phosphoric acid in methanol; gradient 40 to 98% B over 9.5 min; detector wavelength 250 nm; 25 µL injection volume. HPLC Method 2 was used to measure assay (anhydrous basis) *via* pre-column derivatization of the spiroepoxide functionality with potassium pyrimidine-2-thiolate: HPLC Assay (anhydrous basis) 191 mg SDX-7539 / g SDX-7320; (~ 19 wt%); 0.45 mmol SDX-7539 / g SDX-7320;

GPC: The molecular weight distribution was measured by gel permeation chromatography (GPC) on an Agilent 1260 Series instrument or equivalent equipped with refractive index detector using a Styragel HR4 column, 300 x 7.8 mm x 5 µm and elution with hexafluoroisopropanol (HFIP) containing 1 g/L sodium trifluoroacetate at 40^o^C; flow rate 1.0 mL/min; injection volume 20 μL. A calibration curve was established using poly(methyl methacrylate) (PMMA) standards as detected by differential refractometry: M_w_ 29 kDa; M_n_ 19.5 kDa; M_z_ 42 kDa; M_p_ 26 kDa; PDI 1.49;

FTIR: The FTIR-ATR spectrum was obtained using a Perkin-Elmer Spectrum One FTIR spectrophotometer equipped with diamond internal reflection element for ATR, with acquisition range between 3800 – 650 cm^-1^, using 64 scans, with 1 cm^-1^ resolution: ν_max_ (cm^-1^) 3342 (br), 2968, 2930, 1637 (s), 1521 (s), 1456;

Dynamic Light Scattering: Z-Average size and polydispersity index (PDI) were determined by dynamic light scattering (DLS) using a Malvern ZetaSizer ZS90 using the following instrumental parameters: refractive index 1.35; temperature 25^o^C; viscosity 0.89 cP; equilibration time 120 s; 0.5 mL polystyrene cells; 11 cycles; dissolution in 5% aqueous D-mannitol at 20 mg/mL; DLS data recorded after 24 hours of storage at ambient conditions: monomodal; Z-Ave 10.3 nm; PDI 0.23;

NMR: NMR spectra were acquired on a Varian Inova spectrometer at 500 MHz for ^1^H NMR and 125 MHz for ^13^C NMR referenced to the deuterated solvent stated. ^1^H and ^13^C Resonance assignments are summarized in Table S1 according to the numbering shown in Figure S1.

**Figure S1. SDX-7320 atom numbering for NMR assignments**

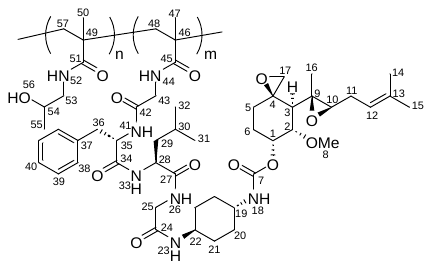

**Table S1.** **^1^H and ^13^C Resonance assignments**

| **Atom** | **^1^H δ (ppm), multiplicity *J* (Hz)**  **(500 MHz, *d*_4_-CH_3_OH)** | **^13^C δ (ppm)**  **(125 MHz, *d*_4_-CH_3_OH)** |
| --- | --- | --- |
| C^1^H | 5.45, a-s | 68.06 |
| C^2^H | 3.66, m | 80.84 |
| C^3^H | 1.94, m | 49.32 |
| C^4^ | - | 60.69 |
| C^5^H_2_ | 2.08, m; 1.05, m | 30.24 |
| C^6^H_2_ | 1.91, m; 1.80, m | 26.89 |
| C^7^=O | - | 157.67 |
| OC^8^H_3_ | 3.42, m | 56.87 |
| C^9^ | - | 60.46 |
| C^10^H | 2.61, t (7.5) | 62.37 |
| C^11^H_2_ | 2.32, m; 2.21, m | 28.26 |
| C^12^H | 5.22, t (7.5) | 119.78 |
| C^13^ | - | 135.89 |
| C^14^H_3_ | 1.75, m | 25.92 |
| C^15^H_3_ | 1.67, m | 18.10 |
| C^16^H_3_ | 1.19, m | 14.29 |
| C^17^H_2_ | 2.56, s; 2.95, m | 51.63 |
| N^18^H | - | - |
| C^19^H | 3.38, m | 50.62 |
| C^20^H_2_ | 1.40, m; 1.92, m | 32.24 |
| C^21^H_2_ | 1.95, m; 1.33, m | 32.48; 32.64 |
| C^22^H | 3.65, m | 49.37 |
| N^23^H | - | - |
| C^24^=O | - | 170.46 |
| C^25^H_2_ | 3.86, m; 3.75, m | 43.68 |
| N^26^H | - | - |
| C^27^=O | - | 174.84 |
| C^28^H | 4.24, m | 54.10 |
| C^29^H_2_ | 1.65, m | 41.07 |
| C^30^H | 1.64, m | 25.74 |
| C^31^H_3_ | 0.94, m | 23.55 |
| C^32^H_3_ | 0.92, m | 22.31 |
| N^33^H | - | - |
| C^34^=O | - | 173.80 |
| C^35^H | 4.64, m | 56.87 |
| C^36^H_2_ | 3.19, m; 2.94, m | 38.56 |
| C^37^ | - | 138.31 |
| C^38^H | 7.29, m | 129.59 |
| C^39^H | 7.27, m | 130.35 |
| C^40^H | 7.22, m | 127.85 |
| N^41^H | - | - |
| C^42^=O | - | 171.71 |
| C^43^H_2_ | 3.78, m; 3.62, m | 43.68 |
| N^44^H | - | - |
| C^45^=O | - | 179.64 |
| C^46^ | - | 46.81 |
| C^47^H_3_ | 1.01, m | 19.38 |
| C^48^H_2_ | 1.88, m | 53.34 |
| C^49^ | - | 46.41 |
| C^50^H_3_ | 1.01, m | 18.25 |
| C^51^=O | - | 180.03 |
| N^52^H | - | - |
| C^53^H_2_ | 3.16, m; 3.00, m | 48.86 |
| C^54^H | 3.87, m | 66.78; 66.92 |
| C^55^H_3_ | 1.14, m | 21.58; 21.37 |
| O^56^H | - | - |
| C^57^H_2_ | 1.74, m | 55.97 |

s = singlet; m = multiplet; a = apparent

**(3*R*,4*S*,5*S*,6*R*)-5-Methoxy-4-((2*R*,3*R*)-2-methyl-3-(3-methylbut-2-en-1-yl)oxiran-2-yl)-1-oxaspiro[2.5]octan-6-yl ((1*R*,4*r*)-4-acetamidocyclohexyl)carbamate (SDX-9246):**

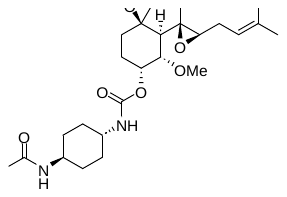

To a mixture of (3*R*,4*S*,5*S*,6*R*)-5-methoxy-4-((2*R*,3*R*)-2-methyl-3-(3-methylbut-2-en-1-yl)oxiran-2-yl)-1-oxaspiro[2.5]octan-6-yl ((1*R*,4*r*)-4-aminocyclohexyl)carbamate (700 mg, 1.44 mmol) in dichloromethane (10 mL) was added acetic anhydride (0.47 mL, 4.96 mmol) and triethylamine (0.69 mL, 4.96 mmol) at 0^o^C. The reaction was stirred for 3 h at room temperature. The reaction mixture was diluted with dichloromethane (300 mL) and washed with water (150 mL) and brine (150 mL). The organic layer was dried over anhydrous sodium sulfate, the solvent was evaporated and the crude residue was purified by flash chromatography (SiO_2_: CH_2_Cl_2_ = 100% to 10% MeOH in CH_2_Cl_2_) to obtain (3*R*,4*S*,5*S*,6*R*)-5-methoxy-4-((2*R*,3*R*)-2-methyl-3-(3-methylbut-2-en-1-yl)oxiran-2-yl)-1-oxaspiro[2.5]octan-6-yl ((1*R*,4*r*)-4-acetamidocyclohexyl)carbamate as a white foam (650 mg, 84%): *m/z* (APCI^+^) 465.2 [M + H]^+^ 100%; ^1^H NMR (400 MHz, *d*_6_-DMSO) *δ* 7.64 (d, *J* 8 Hz, 1H), 6.94 (d, *J* 8 Hz, 1H), 5.23 (br-s, 1H), 5.15 (t, *J* 8 Hz, 1H), 3.49 (dd, *J* 8 Hz, 4 Hz, 1H), 3.39 (br-s, 1H), 3.24 (s, 3H), 2.80 (d, *J* 4 Hz, 1H), 2.53 (d, *J* 4 Hz, 1H), 2.17 (m, 2H), 1.89-1.77 (m, 10H), 1.7 (s, 3H), 1.6 (s, 3H), 1.2-1.1 (m, 7H), 1.1 (s, 3H), 1.04-0.99 (m, 1H).

**(3*R*,4*S*,5*S*,6*R*)-5-Methoxy-4-((2*R*,3*R*)-2-methyl-3-(3-methylbut-2-en-1-yl)oxiran-2-yl)-1-oxaspiro[2.5]octan-6-yl ((1*r*,4*R*)-4-(17-oxo-21-((3a*S*,4*S*,6a*R*)-2-oxohexahydro-1*H*-thieno[3,4-*d*]imidazol-4-yl)-4,7,10,13-tetraoxa-16-azahenicosanamido)cyclohexyl)carbamate (SDX-9280):**

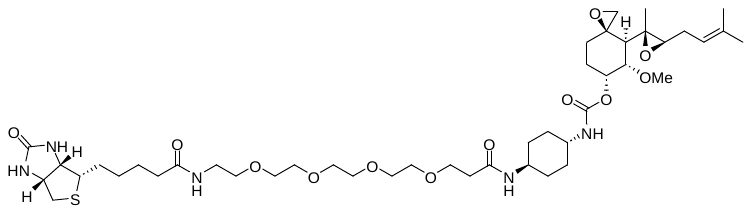

To a mixture of (3*R*,4*S*,5*S*,6*R*)-5-methoxy-4-((2*R*,3*R*)-2-methyl-3-(3-methylbut-2-en-1-yl)oxiran-2-yl)-1-oxaspiro[2.5]octan-6-yl ((1*r*,4*R*)-4-aminocyclohexyl)carbamate (870 mg, 2.06 mmol) in *N,N*-dimethylformamide (10 mL) was added NHS-PEG_4_-Biotin (1.21 g, 2.06 mmol) and *N*,*N*-diisopropylethylamine (1.1 mL, 6.18 mmol) at 0^o^C. The reaction was stirred at room temperature for 15 h. The reaction mixture was concentrated, and the residue was dissolved in water (50 mL) and extracted with ethyl acetate (3 x 50 mL). The aqueous layer was freeze-dried, and the crude compound was purified by reverse phase chromatography (C_18_: H_2_O/CH_3_CN (0.1% trifluoroacetic acid)). Product-containing fractions were freeze-dried to give the isolated product as a white foam (940 mg, 51%): *m/z* (APCI^+^) 896.5 [M + H]^+^ 100%; ^1^H NMR (300 MHz, *d*_6_-DMSO) *δ* 7.84 (t, *J* 5.4 Hz, 1H), 7.72 (t, *J* 7.5 Hz, 1H), 7.01 (d, *J* 7.8 Hz, 1H), 6.42 (s, 1H), 6.36 (s, 1H), 5.27 (br-s, 1H), 5.18 (t, *J* 7.8 Hz, 1H), 4.32-4.27 (m, 1H), 4.32-4.27 (m, 1H), 3.56 (t, *J* 6.3 Hz, 2H), 3.49-3.44 (m, 12H), 3.38 (t, *J* 6.1 Hz, 2H), 3.27 (s, 3H), 3.2-3.07 (m, 4H), 2.80 (dd, *J* 12.3, 5.1 Hz, 2H), 2.56 (dd, *J* 6.3, 5.7 Hz, 2H), 2.26 (t, *J* 6.3 Hz, 2H), 2.17 (m, 2H), 2.05 (t, *J* 7.8 Hz, 2H), 1.89-1.74 (m, 9H), 1.7 (s, 3H), 1.6 (s, 3H), 1.51-1.42 (m, 4H), 1.3-1.1 (m, 8H), 1.1 (s, 3H), 1.04-0.99 (m, 1H).

**Diastereomeric mixture obtained from acidic hydrolysis of fumagill-6-yl *N*-(*trans*-4-aminocyclohexyl)carbamate (SDX-9178):**

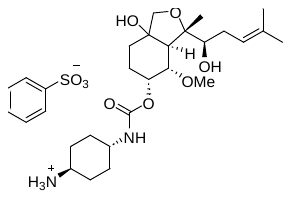

To a mixture of fumagill-6-yl *N*-(*trans*-4-aminocyclohexyl)carbamate (800 mg, 1.9 mmol) in acetonitrile/water (1:1 v/v, 14 mL) was added trifluoroacetic acid (2.2 mL) at 0^o^C. The reaction was stirred at room temperature for 15 h. The reaction mixture was evaporated and the crude residue was purified by reverse phase chromatography (C_18_: H_2_O/CH_3_CN (0.1% trifluoroacetic acid)). Product-containing fractions were freeze-dried to give the isolated product as a mixture of diastereomers in its trifluoroacetate salt form (240 mg): *m/z* (APCI^+^) 441.3 [M + H]^+^ 100%; ^1^H NMR (300 MHz, *d*_6_-DMSO) *δ* 7.77 (br-s, 5H), 7.13 (d, *J* 7.2 Hz, 1H), 6.99 (d, *J* 7.5 Hz, 1H), 5.27-5.20 (m, 2H), 5.10 (br-s, 1H), 3.69 (d, *J* 8.4 Hz, 1H), 3.57-3.32 (m, 3H), 3.27 (s, 3H), 2.93 (br-s, 1H), 2.23 (m, 2H), 1.89-1.77 (m, 9H), 1.70 (s, 3H), 1.60 (s, 3H), 1.40-1.10 (m, 10H). To a solution of the trifluoroacetate salt (960 mg, 1.73 mmol) in water (25 mL) was added aqueous sodium hydroxide (0.1 N, 20 mL) and the mixture was stirred for 10 min at room temperature. The mixture was extracted with ethyl acetate (3 x 100 mL) and washed with water (150 mL) and brine (150 mL). The organic layer was dried over anhydrous sodium sulfate and evaporated to provide the product as a gum in its free base form (520 mg). To a mixture of the free base (520 mg, 1.18 mmol) in 1-butanol (4 mL) was added a solution of benzenesulfonic acid (187 mg, 1.18 mmol) in methyl *tert*-butyl ether (10 mL). The mixture was cooled to 5^o^C for 15 h. The resulting white precipitate was filtered, washed with methyl *tert*-butyl ether (4 mL) and dried to provide the product as a mixture of diastereomers in its benzenesulfonic acid salt form as an off-white solid (400 mg, 57% from the free base): *m/z* (APCI^+^) 441.3 [M + H]^+^ 100%; ^1^H NMR (300 MHz, *d*_6_-DMSO) *δ* 7.72 (br-s, 2H), 7.62-7.57 (m, 2H), 7.36-7.27 (m, 3H), 5.23-5.15 (m, 2H), 5.10 (br-s, 1H), 3.72 (m, 1H), 3.57-3.32 (m, 6H), 3.27 (s, 3H), 3.20-3.07 (m, 3H), 2.94 (m, 1H), 1.89-1.64 (m, 12H), 1.39-1.04 (m, 10H).

**Enzyme-mediated metabolism of SDX-7320 *in vitro***

Cathepsin isoforms and human neutrophil elastase (HNE) were tested for their ability to catalyze the release of SDX-7539 from SDX-7320 *in vitro*. All enzymes were diluted to 0.021 U enzyme/mL. The activity of human cathepsins A, B, C, D, E, F, G, H, K, L, O, S, V, W, X/Z/P and mouse cathepsin L, as well as HNE was measured at 37^o^C with shaking (1600 rpm) in assay buffer (13.3 mM sodium acetate, pH 5.0, 1.5 mM EDTA, 3.0 mM DTT) containing 0.5 mg/mL SDX-7320, 200 ng/mL of deuterated SDX-7539 (as an internal standard) although the cathepsin D assay did not contain EDTA or DTT. Reactions were halted by addition of 0.6 volumes of methanol, followed by freezing at -20^o^C for 15 minutes. After centrifugation for 4 minutes (13,000 rpm), a portion of the supernatant from each reaction was transferred to a glass auto-sampler vial and SDX-7539 was measured by LC/MS/MS using a Sciex API 6500 and MRM detection (LLOQ = 48 ng/mL, ULOQ = 200,000 ng/mL). Separate experiments were conducted using cathepsin L to first activate cathepsins A, C and H (enzymes were combined and incubated on ice for 15 minutes in sodium acetate buffer as above with EDTA and DTT) prior to incubation with SDX-7320. In these studies, cathepsin L alone was also tested. Samples were analyzed for SDX-7539 at 1, 2, 4, 6, or 8 hours after addition of SDX-7320 to the enzyme solutions.

**Expression of MetAP2 and MetAP2-SDX-7539 co-crystal structure**

Human MetAP2 108-478 (Uniprot ID: P50579) was expressed using a baculovirus–insect cell system with an N-terminal 6His tag. Protein was purified using Ni IMAC, TEV cleavage, subtractive Ni IMAC and size-exclusion chromatography. Purified protein was concentrated to 22 mg/mL before performing crystal screening in the presence of 1.0 mM MnCl_2_ and 5.0 mM SDX-9402. Protein and compound were incubated for 1 hour prior to setting up crystallization plates. Crystals were obtained within two days in 0.2M ammonium sulfate, 0.1M BIS-TRIS pH 5.5, 25% w/v polyethylene glycol 3,350. MetAP2-SDX-7539 co-crystals were cryoprotected with reservoir buffer supplemented with 20% glycerol before flash-cooling in liquid nitrogen.

X-ray diffraction data from crystals of the MetAP2-SDX-7539 complex were collected at the Soleil synchrotron facility (Saint-Aubin, France) under cryogenic conditions. Details of the data reduction and refinement are provided in Table S5. The final refined structure was deposited in the PDB under code 8OXG.

X-ray diffraction data were processed using AIMLESS. The phase information necessary to determine and analyze the structure was obtained by molecular replacement using PHASER (6). The previously solved structure of MetAP2 (PDB code: 5D6F) was used as the initial search model. Subsequent model building and refinement was performed according to standard protocols using CCP4 (7) and COOT (8) software. The ligand parameterization and the generation of the corresponding library files was carried out with ACEDRG (CCP4). Validation of the structure was achieved using programs MolProbity (9) and PDB validation (10). Structural images and superpositions were generated using Pymol (10).

***In vitro* competition binding to MetAP2**

The sandwich ELISA method was based on a previously described method (11), in which human recombinant MetAP2 was captured on a streptavidin-coated plate with a biotinylated MetAP2 inhibitor (SDX-9280). A standard curve was generated by first incubating a range of MetAP2 concentrations with biotinylated SDX-9280 in a dilution plate followed by transfer into a streptavidin coated-96 well ELISA plate. After a one-hour incubation at room temperature, the unbound portion was removed by washing the plate with assay buffer (PBS Starting Block). Bound MetAP2 was detected using a rabbit monoclonal anti-MetAP2 antibody followed by horse radish peroxidase (HRP)-conjugated goat anti-rabbit antibody and the HRP substrate 3,3', 5,5;-tetramethylbenzidine (TMB). The reaction was stopped with 2N H_2_SO_4_ and the optical density was measured at 450 nm.

The experimental method to measure relative inhibition of MetAP2 was developed using the above ELISA method and tested the following variables: temperature during incubation of MetAP2 with biotinylated SDX-9280, time of incubation (1 hour vs. 2 hours), the order of reagent addition to the ELISA plate and concentration of biotinylated SDX-9280, tested at 1.0, 0.5, 0.25 and 0.03 µM, with different MetAP2 standard curve ranges. Streptavidin high binding capacity coated plates that were utilized to perform this assay have a binding capacity of ~125 pmol biotin per well. MetAP2 concentrations were optimized for both the standard curve range and the test article compounds. The final conditions were as follows: 0.25 µM of biotinylated SDX-9280 as the capture reagent, assay curve range of 6 to 150 ng/mL of MetAP2, and a one-hour incubation of MetAP2 and SDX-9280 at room temperature plus varying concentrations of test compounds. Compounds (i.e., potential MetAP2 competitors/inhibitors) were tested at various concentrations in the presence of 100 ng/mL MetAP2 and 0.25 mM of SDX-9280.

**MetAP1 enzyme assay**

Human MetAP1 and DPPIV were purchased from R&D Systems. Human MetAP1 (4 μg/mL), CD26 (1 μg/mL), Met-Gly-Pro-AMC (50 μM) and test compounds (0.01 nM to 1000 nM) were incubated at room temperature (50 mM HEPES, 0.1 mM CoCl_2_, 100 mM NaCl, pH 7.5) and fluorescence (excitation at 350 nm, emission at 440 nm) was measured at the indicated times.

**Selectivity screening**

SDX-9402 (benzenesulfonic acid salt form of SDX-7539) was tested at concentrations of 1 x 10^-5^ and 1 x 10^-6^ M in binding, enzyme, and transporter assays (Cerep). Binding of SDX-9402 to the receptors/transporters tested (N=38) or inhibition of enzymes tested (N=6) was calculated as a % inhibition of the binding of a radioactively labeled ligand specific for each target or as a % inhibition of control enzyme activity, respectively. Results showing greater than 50% inhibition or stimulation with SDX-9402 were considered to represent significant effects.

**Pharmacokinetics and CNS exposure of SDX-7539 and TNP-470 in rats:**

The pharmacokinetics and CNS exposure of small-molecule MetAP2 inhibitors was assessed in male Sprague-Dawley rats (n=3/group) following intravenous (IV) doses of SDX-7320, SDX-7539, and TNP-470. The dosing solutions were prepared in PBS. TNP-470 was dissolved in 10% ethanol/90% PBS at 6.0 mg/mL. SDX-7539 solution was filtered in a 0.22 μm filter prior to dosing. At selected time points following bolus IV dosing, blood was collected from each animal via saphenous vein puncture at a volume of 300 µL per time point. Fourteen days after the initial dose the animals were euthanized *via* CO_2_ inhalation and the maximum volume of blood collected via cardiac puncture. Blood samples were centrifuged at 4^o^C at 8,000 RPM for 5 minutes. For assessment of CNS penetration of small molecules, a separate cohort of rats (n=3/group) were euthanized *via* CO_2_ inhalation 10 minutes (small molecules) or 120 minutes after IV dosing (SDX-7320), and blood was collected via cardiac puncture (from which plasma was prepared) and one minute later CSF was collected from each rat then snap frozen. Decanted plasma was stored at -80^o^C.

**Bioanalytical procedures in support of rat pharmacokinetics and CNS exposure**

SDX-7539 and TNP-470 in rat plasma and CSF were measured by LC/MS/MS using either an Agilent 6410 mass spectrometer coupled with an Agilent 1200 HPLC and a CTC PAL chilled auto-sampler, all controlled by MassHunter software (Agilent), or an ABI2000 mass spectrometer coupled with an Agilent 1100 HPLC and a CTC PAL chilled auto-sampler, all controlled by Analyst software (ABI). After separation on a C18 reverse phase HPLC column (Agilent, Waters, or equivalent) using an acetonitrile-water gradient system, peaks were analyzed by mass spectrometry (MS) using ESI ionization in MRM mode.

**Assessment of neurobehavioral effects of SDX-7320 in rats**

Clinical observations were conducted following each FOB examination. FOB evaluations were conducted pre-dose (Day -1) and at 12- and 36-hours post-dose. Dosing (3, 10, or 30 mg/kg, IV) was conducted during the dark cycle in order to ensure all FOB evaluations occurred consistently during the light cycle. The 12-hour interval was initially selected based on the expected T_max_; although the observed toxicokinetics subsequently indicated T_max_ was earlier (4 hours, Figure S4), it was estimated that 47% of SDX-7539 C_max_ and 60% of SDX-7539 AUC was still characterized at 12 hours without any neurobehavioral findings. Results of FOB evaluations are in Table S7. Behavioural components of the FOB were as previously described (12,13). Forelimb and hindlimb grip strength was assessed as described (14). The latency to a nociceptive (thermal) response when the subject was placed on a heated (52 ± 1°C) surface, was measured as described (15). Statistical analysis was conducted using ANOVA with Dunnett’s multiple comparisons (vehicle vs treatment groups).

**A549 Xenograft model**

Female Athymic Nude mice (Hsd:Athymic Nude-Fox-n1^nu^) were supplied by Harlan (Indianapolis, IN). Mice were received at 4-5 weeks of age. All mice were acclimated for one week prior to handling. The mice were housed in microisolator cages (Lab Products, Seaford, DE) and maintained under specific pathogen-free conditions. The mice were fed Tekland Global Diet® 2920x irradiated laboratory animal diet (Harlan; Indianapolis, IN) and provided autoclaved water ad libitum. A549 cells were maintained in RPMI-1640 (Lonza; Walkersville, MD) supplemented with 10% Fetal Bovine Serum (FBS; Seradigm; Radnor, PA) and housed in a humidified, 5% CO_2_ atmosphere at 37°C.

SDX-7539 and SDX-7320 formulations were prepared as follows: 0.9% NaCl solution (saline; B. Braun Medical, Inc., Irvine, CA, ref# L8000) was added to the pre-weighed drug vials, the solutions were swirled for approximately 1 minute and held at room temperature for approximately one hour prior to dosing. SDX-7539 was formulated at concentrations of 3.7 mg/mL to dose 37 mg/kg, respectively, at a 10 mL/kg dose volume. SDX-7320 was formulated at 6 mg/mL to dose 60 mg/kg at a 10 mL/kg dose volume. The 6 mg/mL stock of SDX-7320 was diluted with saline to a concentration of 0.6 mg/mL to dose 6 mg/kg, at a 10 mL/kg dose volume. All SDX drugs were formulated fresh each day of dosing. TNP-470 was dissolved in 100% EtOH (ethanol; EMD Chemicals Inc.; Gibbstown, NJ, cat# EX0276-1), then diluted in PBS (phosphate buffered saline; Hyclone; Logan, UT, cat# SH30256.02) to a concentration of 3 mg/mL (at a final formulation of 10% EtOH; 90% PBS) to dose 30 mg/kg at a 10 mL/kg dose volume. TNP-470 was formulated fresh each day of dosing.

**Histological analysis of EO771 tumors**

Tumor tissue (fixed in buffered formalin for 24 h, then transferred to 70% EtOH) was embedded in paraffin, sectioned and mounted on glass slides. Staining for Ki67 (Abcam ab16667) and CD31 (Abcam ab28364) was conducted using standard techniques (Wax-It Histology Services).

**B16-F10 Lung metastasis model**

Female C57/Bl6 mice (CrTac:C57BL/6NTac) (Taconic, Germantown, NY) were received at four weeks of age and acclimated for seven days prior to handling. The mice were housed in microisolator cages (Lab Products, Seaford, DE) and maintained under specific pathogen-free conditions. The mice were fed PicoLab® irradiated mouse chow (Lab Diet, Richmond, IN) and autoclaved water was freely available.

Mice were inoculated with 0.2 mL of sterile saline containing a suspension of B16-F10 tumor cells (approximately 1 x 10^5^ cells/mouse) IV in the lateral tail vein. One day following inoculation, mice were randomized by body weight into groups of eight mice/group. Body weights were recorded when dosing was initiated, and every other day thereafter. Gross observations were also recorded daily.

TNP-470 and SDX-7320 were stored at -20ºC until use. A stock solution of TNP-470 was made at 100 mg/mL in 100% EtOH), then diluted in PBS (Hyclone; Logan, UT) to a concentration of 3 mg/mL to dose 30 mg/kg at a 10 mL/kg dose volume. SDX-7320, was formulated in 0.9% NaCl solution (saline) at a concentration of 2.5 mg/mL to dose 25 mg/kg, respectively, at a 10 mL/kg dose volume. The preparation of SDX-7320 were as follows: saline was added to vials containing the lyophilized drug, the solution was swirled for approximately 1 minute, and the solution was kept at room temperature for one hour prior to dosing. Treatment was initiated 24 hours after tumor cells had been administered.

**Growth** **of MDA-MB-231 tumors in chick embryos**

Fertilized White Leghorn eggs were incubated at 37.5°C with 50% relative humidity and processed as described (16). Nine days post-fertilization (E9) the chorioallantoic membrane (CAM) was dropped by drilling a small hole through the eggshell into the air sac, and a 1 cm² window was cut in the eggshell above the CAM. MDA-MB-231 tumor cells (ATCC, Manassas, VA) were cultivated in DMEM medium with 10% FBS and 1% penicillin/streptomycin. On day E9, cells were detached and an inoculum of 1x10^6^ cells was added onto the CAM of each egg and then eggs were randomized into groups (n = 35–38 eggs/group). Treatment with test agents was initiated on day 11 post-fertilization (E11). Vehicle (0.5% mannitol in water, Q4D, two doses; 0.25% DMSO in PBS, QOD, four doses), paclitaxel (2.1 μg in 0.1 mL 0.25% DMSO in PBS), SDX-7320 (15 or 90 μg in 0.5% aqueous mannitol).

Quantitation of angiogenesis was conducted on day E16 using an image of the upper CAM (with tumor) captured using a digital camera. The number of blood vessels that reach the tumor was counted in triplicate to evaluate tumor angiogenesis (n=8 eggs/group). Tumor weight was measured on day E18, after the upper portion of the CAM (with tumor) was removed, washed in PBS and then transferred into 4% paraformaldehyde for 48 h. Tumors were then cut away from normal CAM tissue and weighed (14 to 24 tumors from each group, depending on survival at day E18). Metastatic invasion of MDA-MB-231 tumor cells into the embryo (lower CAM) was measured on day E18 by first dissecting a portion of the lower CAM (1 cm^2^, n=8/group). Genomic DNA was extracted from the tissue (NuceloMag 96 Tissue, Macherey-Nagel, Germany), and analyzed by qPCR with specific primers for human Alu sequences. Calculation of Cq for each sample, mean Cq and relative amounts of metastases for each group were determined using Bio-Rad® CFX Maestro software.

**Figure S2. *In vitro* Competition binding to MetAP2.** The MetAP2 binding assay relied upon a sandwich ELISA in which a standard curve was created by first combining 0.25 µM of biotinylated SDX-7539 (SDX-9280) with varying concentrations of human recombinant MetAP2. An aliquot of this reaction mixture was transferred to a 96-well streptavidin plate, and incubated at room temperature for one hour. The plate was then washed and an anti-MetAP2 antibody was added, plus an HRP-conjugated goat anti-rabbit antibody and TMB. The reaction was stopped with 2N H_2_SO_4_ and the optical density was measured at 450 nanometers. The assay was linear from 6 to 150 ng/mL of MetAP-2. Inhibition samples for competition binding were tested at 100 ng/mL MetAP-2, 0.25 µM of biotinylated SDX-7539 (SDX-9280) and varying concentrations of test compounds.

A. **B.**

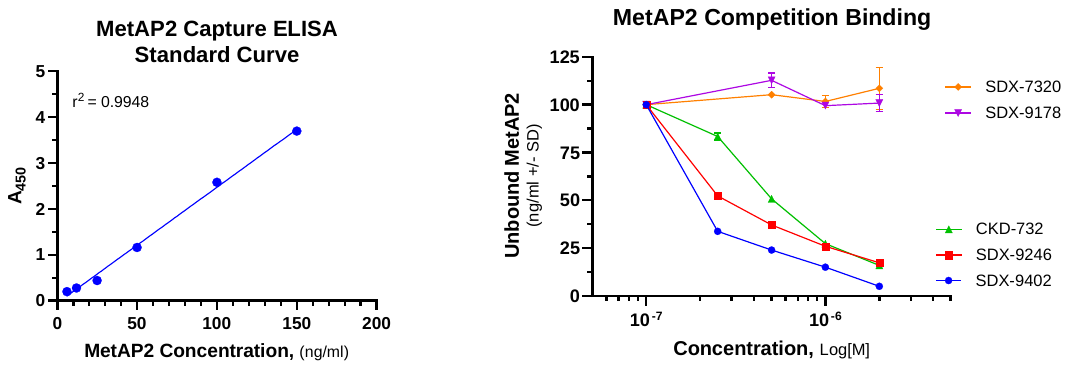

**Figure S3. Superposition of crystal structures MetAP2-SDX-7539 and MetAP2-TNP-470 (PDB code 1B6A)**

Protein conformation and bound compound position were highly similar between MetAP2-SDX-7539 (blue and pink) and MetAP2-TNP-470 (purple and green). An RMSD of 0.245Å was calculated across all atoms.

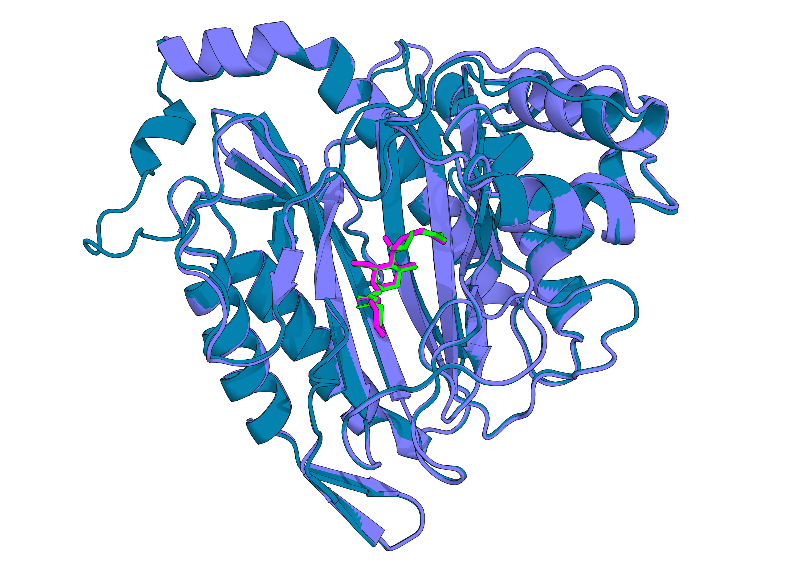

**Figure S4. MetAP1 enzyme assay.** Enzyme activity was measured at room temperature using a coupled assay system (R&D Systems) containing the fluorogenic substrates Met-Gly-Pro-AMC, 4 μg/mL recombinant human MetAP1, 1 μg/mL DPPIV, and 1 μM Met-Gly-Pro-AMC in 50 mM HEPES, 0.1 mM CoCl_2_, 100 mM NaCl, pH 7.5. Test compounds were diluted in DMSO, then diluted in 96-well plates containing enzymes, followed by addition of the fluorogenic substrate. Fluorescence was measured over time using excitation and emission wavelengths of 350 nm and 440 nm, respectively.

A. **B.**

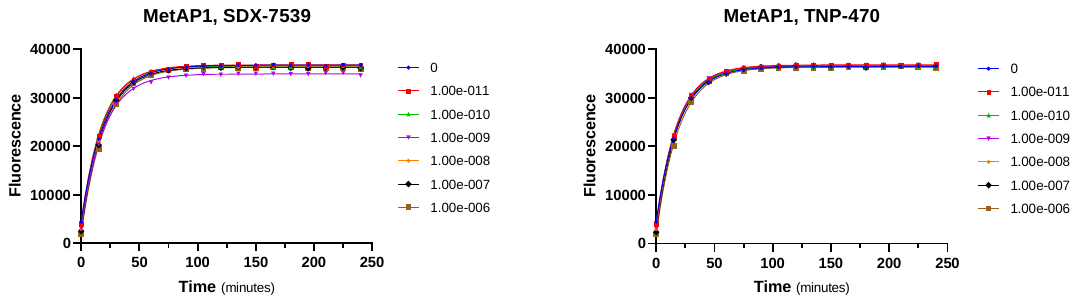

**Figure S5. Pharmacokinetics of SDX-7539 in rats.** The test agents SDX-7320 (dosing solution prepared in PBS at 40 mg/mL) and SDX-7539 (dosing solution prepared in PBS at 7.4 mg/mL) were dosed intravenously at time zero (n=3/group), and blood (0.3 mL/time point) was collected from each animal via saphenous vein puncture. Blood was collected into K_2_EDTA tubes for plasma separation. Blood samples were centrifuged at 4^0^C at 8,000 RPM for 5 minutes and the supernatant plasma was stored at -70^0^C until analysis.

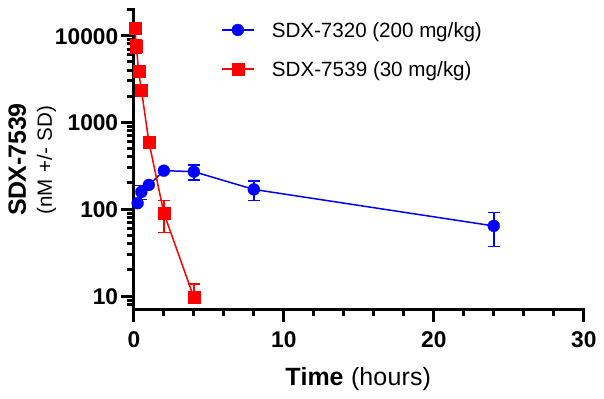

**Figure S6. Principle component analysis (PCA) of BT-474 RNASeq Data.** Principal component analysis (PCA) was performed on RNASeq data using the PCA function within the python package scikit-learn. Normalized gene expression was used as input to PCA to find the first 10 components of global gene expression.

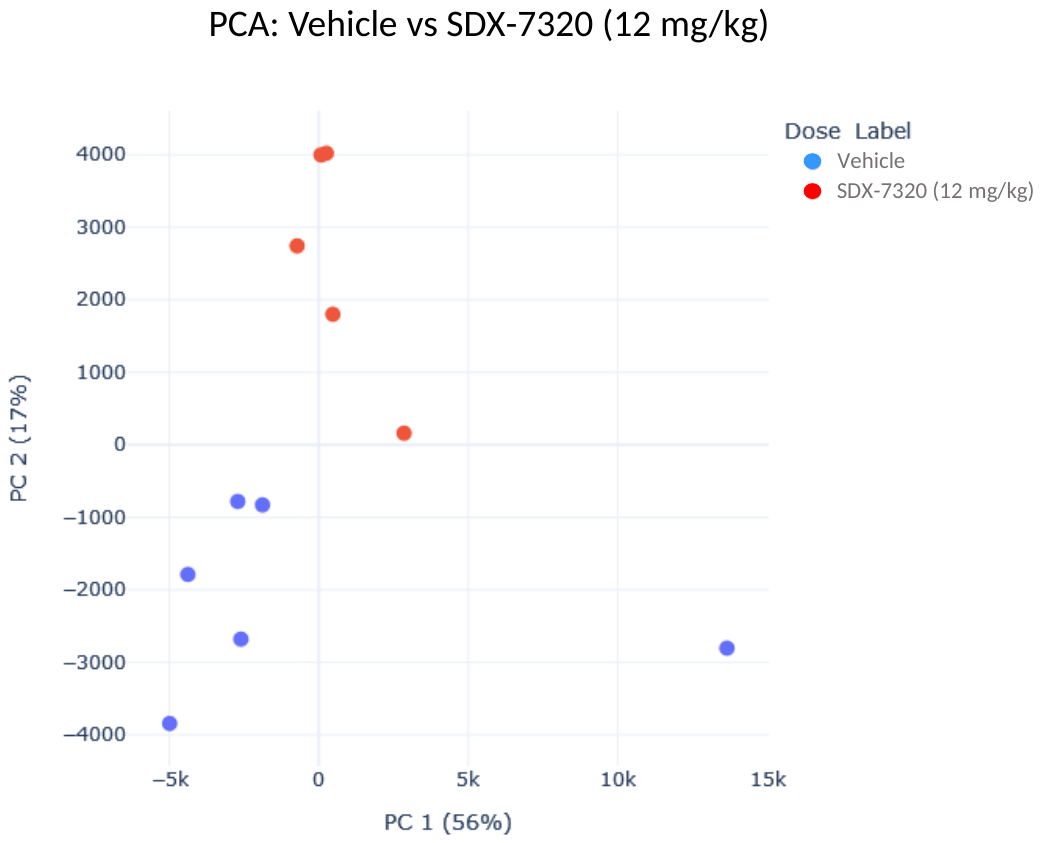

**Table S2. Commercial sources of cathepsins**

| Cathepsin | Subtype | Description | Vendor | CODE |
| --- | --- | --- | --- | --- |
| A | Serine | HUMAN, Recombinant | R&D Systems | 1049-SE-010 |
| B | Cysteine | HUMAN, Recombinant | R&D Systems | 953-CY-010 |
| C | Cysteine | HUMAN, Recombinant | R&D Systems | 1071-CY-010 |
| D | Aspartic | HUMAN, Recombinant | R&D Systems | 1014-AS-010 |
| E | Aspartic | HUMAN, Recombinant | R&D Systems | 1294-AS-010 |
| F | Cysteine | HUMAN, Recombinant | ABCAM | AB198436 |
| G | Serine | HUMAN Neutrophils, Purified | ENZO Life Sciences | BML-SE283-0100 |
| H | Cysteine | HUMAN, Recombinant | R&D Systems | 7516-CY-010 |
| K | Cysteine | HUMAN, Recombinant | ENZO Life Sciences | BML-SE553-010 |
| L | Cysteine | HUMAN, Recombinant | R&D Systems | 952-CY-010 |
| O | Cysteine | HUMAN, Recombinant | CREATIVE Biomart | CTSO-1103H |
| S | Cysteine | HUMAN, Recombinant | R&D Systems | 1183-CY-010 |
| V | Cysteine | HUMAN, Recombinant | R&D Systems | 1080-CY-010 |
| W | Cysteine | HUMAN, Recombinant | CREATIVE Biomart | CTSW-8183H |
| X/Z/P | Cysteine | HUMAN, Recombinant | R&D Systems | 934-CY-010 |
| HNE | Serine | HUMAN, Purified | R&D Systems | 9167-SE-020 |
| mL | Cysteine | MOUSE, Recombinant | R&D Systems | 2336-CY-010 |
| IU = μmol/min | |  |  |  |

**Table S3. Measurement of off-target binding of SDX-9402.** Binding of SDX-9402 (benzenesulfonic acid salt form of SDX-7539, tested at concentrations of 1 x 10^-5^ and 1 x 10^-6^ M) to the receptors/transporters tested (N=38) was calculated as % inhibition of the binding of a radioactively labeled ligand specific for each target. Results showing greater than 50% inhibition or stimulation with SDX-9402 were considered to represent significant effects.

|  |  |  | **% Inhibition of Specific Binding** | | |
| --- | --- | --- | --- | --- | --- |
| **Receptor/Transporter** | **Compound** | **Test Concentration (mol/L)** | **1st** | **2nd** | **Mean** |
| A2A |  | (agonist radioligand) |  |  |  |
|  | SDX-9402 | 1.00E-06 | 0.3 | 6.4 | 3.3 |
|  | SDX-9402 | 1.00E-05 | 2.7 | -4.4 | -0.8 |
| α1A |  | (antagonist radioligand) |  |  |  |
|  | SDX-9402 | 1.00E-06 | -1.9 | 4.4 | 1.3 |
|  | SDX-9402 | 1.00E-05 | 15.7 | 10.9 | 13.3 |
| α2A |  | (antagonist radioligand) |  |  |  |
|  | SDX-9402 | 1.00E-06 | 2.3 | 2.5 | 2.4 |
|  | SDX-9402 | 1.00E-05 | 13.6 | 17.8 | 15.7 |
| β1 |  | (agonist radioligand) |  |  |  |
|  | SDX-9402 | 1.00E-06 | 9.4 | 3.9 | 6.7 |
|  | SDX-9402 | 1.00E-05 | 10.6 | 4.2 | 7.4 |
| β2 |  | (agonist radioligand) |  |  |  |
|  | SDX-9402 | 1.00E-06 | 2.5 | 11.1 | 6.8 |
|  | SDX-9402 | 1.00E-05 | 7 | 3.3 | 5.2 |
| Benzodiazepine (central) |  | (agonist radioligand) |  |  |  |
|  | SDX-9402 | 1.00E-06 | -0.2 | 3.3 | 1.6 |
|  | SDX-9402 | 1.00E-05 | 9.4 | 14.2 | 11.8 |
| CB1 |  | (agonist radioligand) |  |  |  |
|  | SDX-9402 | 1.00E-06 | -9.9 | -14.7 | -12.3 |
|  | SDX-9402 | 1.00E-05 | -21.5 | -10.1 | -15.8 |
| CB2 |  | (agonist radioligand) |  |  |  |
|  | SDX-9402 | 1.00E-06 | 23.6 | 4.4 | 14 |
|  | SDX-9402 | 1.00E-05 | -7.3 | -2.3 | -4.8 |
| CCK1 (CCKA) |  | (agonist radioligand) |  |  |  |
|  | SDX-9402 | 1.00E-06 | -39.9 | -29.3 | -34.6 |
|  | SDX-9402 | 1.00E-05 | -18.3 | -7.9 | -13.1 |
| D1 |  | (antagonist radioligand) |  |  |  |
|  | SDX-9402 | 1.00E-06 | 13.5 | 11.3 | 12.4 |
|  | SDX-9402 | 1.00E-05 | 3.6 | 10 | 6.8 |
| D2S |  | (agonist radioligand) |  |  |  |
|  | SDX-9402 | 1.00E-06 | -2.6 | -7.4 | -5 |
|  | SDX-9402 | 1.00E-05 | -21.6 | 4.1 | -8.7 |
| ETA |  | (agonist radioligand) |  |  |  |
|  | SDX-9402 | 1.00E-06 | 15.3 | 14.1 | 14.7 |
|  | SDX-9402 | 1.00E-05 | 2 | 3.3 | 2.7 |
| NMDA |  | (antagonist radioligand) |  |  |  |
|  | SDX-9402 | 1.00E-06 | 0.4 | -6.2 | -2.9 |
|  | SDX-9402 | 1.00E-05 | 6.3 | 0 | 3.1 |
| H1 |  | (antagonist radioligand) |  |  |  |
|  | SDX-9402 | 1.00E-06 | -4.8 | 1.9 | -1.4 |
|  | SDX-9402 | 1.00E-05 | -6.5 | 2.6 | -2 |
| H2 |  | (antagonist radioligand) |  |  |  |
|  | SDX-9402 | 1.00E-06 | 14.5 | 0.8 | 7.6 |
|  | SDX-9402 | 1.00E-05 | -12 | -1.7 | -6.8 |
| MAO-A |  | (antagonist radioligand) |  |  |  |
|  | SDX-9402 | 1.00E-06 | -11.1 | -4.3 | -7.7 |
|  | SDX-9402 | 1.00E-05 | -10.2 | 7.6 | -1.3 |
| M1 |  | (antagonist radioligand) |  |  |  |
|  | SDX-9402 | 1.00E-06 | 2.7 | -7.3 | -2.3 |
|  | SDX-9402 | 1.00E-05 | 5.5 | -0.8 | 2.4 |
| M2 |  | (antagonist radioligand) |  |  |  |
|  | SDX-9402 | 1.00E-06 | -5.6 | -0.8 | -3.2 |
|  | SDX-9402 | 1.00E-05 | 2.8 | 7.4 | 5.1 |
| M3 |  | (antagonist radioligand) |  |  |  |
|  | SDX-9402 | 1.00E-06 | 7 | 0 | 3.5 |
|  | SDX-9402 | 1.00E-05 | -5 | 6.2 | 0.6 |
| N neuronal a4b2 |  | (agonist radioligand) |  |  |  |
|  | SDX-9402 | 1.00E-06 | 3.9 | 8.4 | 6.2 |
|  | SDX-9402 | 1.00E-05 | 9.8 | 2.6 | 6.2 |
| δ2 (DOP) |  | (agonist radioligand) |  |  |  |
|  | SDX-9402 | 1.00E-06 | 3.2 | -5.1 | -1 |
|  | SDX-9402 | 1.00E-05 | 1.4 | -0.9 | 0.3 |
| κ (KOP) |  | (agonist radioligand) |  |  |  |
|  | SDX-9402 | 1.00E-06 | 18.9 | 5.4 | 12.1 |
|  | SDX-9402 | 1.00E-05 | 5.5 | 21.8 | 13.7 |
| μ (MOP) |  | (agonist radioligand) |  |  |  |
|  | SDX-9402 | 1.00E-06 | 1.9 | 0.5 | 1.2 |
|  | SDX-9402 | 1.00E-05 | 11.7 | 5.5 | 8.6 |
| 5-HT1A |  | (agonist radioligand) |  |  |  |
|  | SDX-9402 | 1.00E-06 | 24 | 12.2 | 18.1 |
|  | SDX-9402 | 1.00E-05 | 21.2 | 16.5 | 18.9 |
| 5-HT1B |  | (antagonist radioligand) |  |  |  |
|  | SDX-9402 | 1.00E-06 | -11.6 | 4.8 | -3.4 |
|  | SDX-9402 | 1.00E-05 | -20.7 | -20.1 | -20.4 |
| 5-HT2A |  | (agonist radioligand) |  |  |  |
|  | SDX-9402 | 1.00E-06 | -20.3 | -22.8 | -21.5 |
|  | SDX-9402 | 1.00E-05 | -9.4 | -10.8 | -10.1 |
| 5-HT2B |  | (agonist radioligand) |  |  |  |
|  | SDX-9402 | 1.00E-06 | 2 | -1.6 | 0.2 |
|  | SDX-9402 | 1.00E-05 | -3.2 | -4.7 | -3.9 |
| 5-HT3 |  | (antagonist radioligand) |  |  |  |
|  | SDX-9402 | 1.00E-06 | -7.1 | 6.4 | -0.3 |
|  | SDX-9402 | 1.00E-05 | 2.4 | 16.7 | 9.5 |
| GR |  | (agonist radioligand) |  |  |  |
|  | SDX-9402 | 1.00E-06 | -17.3 | -15.5 | -16.4 |
|  | SDX-9402 | 1.00E-05 | -13.7 | -15 | -14.4 |
| AR |  | (agonist radioligand) |  |  |  |
|  | SDX-9402 | 1.00E-06 | 3.5 | 13.5 | 8.5 |
|  | SDX-9402 | 1.00E-05 | 16.9 | 13.1 | 15 |
| V1a |  | (agonist radioligand) |  |  |  |
|  | SDX-9402 | 1.00E-06 | 0.9 | -8.2 | -3.6 |
|  | SDX-9402 | 1.00E-05 | 3.2 | 8.1 | 5.7 |
| Ca2+ channel dihydropyridine site |  | (antagonist radioligand) |  |  |  |
|  | SDX-9402 | 1.00E-06 | -10.5 | -9.9 | -10.2 |
|  | SDX-9402 | 1.00E-05 | -10.7 | -14.3 | -12.5 |
| hERG |  | (antagonist radioligand) |  |  |  |
|  | SDX-9402 | 1.00E-06 | -16 | 23.4 | 3.7 |
|  | SDX-9402 | 1.00E-05 | 2.1 | -16.5 | -7.2 |
| KV channel |  | (antagonist radioligand) |  |  |  |
|  | SDX-9402 | 1.00E-06 | -9.2 | -3.6 | -6.4 |
|  | SDX-9402 | 1.00E-05 | -3.8 | -1.8 | -2.8 |
| Na+ channel (site 2) |  | (antagonist radioligand) |  |  |  |
|  | SDX-9402 | 1.00E-06 | 22.3 | 9.4 | 15.9 |
|  | SDX-9402 | 1.00E-05 | 19.7 | 39.8 | 29.7 |
| Norepinephrine transporter |  | (antagonist radioligand) |  |  |  |
|  | SDX-9402 | 1.00E-06 | 1.7 | 8.3 | 5 |
|  | SDX-9402 | 1.00E-05 | -0.3 | 9.5 | 4.6 |
| Dopamine transporter |  | (antagonist radioligand) |  |  |  |
|  | SDX-9402 | 1.00E-06 | 0.3 | 3.9 | 2.1 |
|  | SDX-9402 | 1.00E-05 | -0.3 | 14.4 | 7 |
| 5-HT transporter |  | (antagonist radioligand) |  |  |  |
|  | SDX-9402 | 1.00E-06 | 3.8 | 8.4 | 6.1 |
|  | SDX-9402 | 1.00E-05 | 9.1 | 0.1 | 4.6 |

**Table S4. Measurement of off-target enzyme inhibition by SDX-9402.** Inhibition of enzymes (N=6) by SDX-9402 (benzenesulfonic acid salt form of SDX-7539, tested at concentrations of 1 x 10^-5^ and 1 x 10^-6^ M) was calculated as a % inhibition of control enzyme activity. Results showing greater than 50% inhibition or stimulation with SDX-9402 were considered to represent significant effects.

|  |  |  | **% Inhibition of Enzyme Activity** | | |
| --- | --- | --- | --- | --- | --- |
| **Enzyme** | **Compound** | **Test Concentration (mol/L)** | **1st** | **2nd** | **Mean** |
| COX1 |  |  |  |  |  |
|  | SDX-9402 | 1.00E-06 | -16.5 | 19.8 | 1.7 |
|  | SDX-9402 | 1.00E-05 | 17.1 | 3.4 | 10.3 |
| COX2 |  |  |  |  |  |
|  | SDX-9402 | 1.00E-06 | 15.4 | 15 | 15.2 |
|  | SDX-9402 | 1.00E-05 | 15.6 | 13.8 | 14.7 |
| PDE3A |  |  |  |  |  |
|  | SDX-9402 | 1.00E-06 | -0.3 | -1.4 | -0.8 |
|  | SDX-9402 | 1.00E-05 | 0.6 | 1.5 | 1 |
| PDE4D2 |  |  |  |  |  |
|  | SDX-9402 | 1.00E-06 | -3.2 | 6.3 | 1.6 |
|  | SDX-9402 | 1.00E-05 | -1.9 | -1.8 | -1.8 |
| Lck |  |  |  |  |  |
|  | SDX-9402 | 1.00E-06 | 3 | 3.9 | 3.4 |
|  | SDX-9402 | 1.00E-05 | 18.6 | 4.2 | 11.4 |
| Acetylcholinesterase |  |  |  |  |  |
|  | SDX-9402 | 1.00E-06 | 3.9 | 7.3 | 5.6 |
|  | SDX-9402 | 1.00E-05 | 3.2 | 6.6 | 4.9 |

**Table S5. Data collection and refinement statistics for MetAP2-SDX-7539 co-crystal structure**

| **Data Processing** | |
| --- | --- |
| X-ray source | PROXIMA-1 Soleil |
| Collection date | 14/06/2022 |
| Wavelength [Å] | 0.9786 |
| Detector | EIGER-X 16M |
| Temperature [K] | 100 |
| Space group | C121 |
| Cell: a; b; c; [Å] | 119.05 101.13 83.74 |
| α; β; γ; [◦] | 90.00 98.75 90.00 |
| Resolution [Å] | 44.88-1.73 (1.76-1.73)* |
| Unique reflections | 101909 (4621)* |
| Multiplicity | 6.9 (5.7)* |
| Completeness ([%]) | 99.6 (92.3)* |
| Rmerge [%] | 0.089 (1.288)* |
| Rpim [%] | 0.055 (0.866)* |
| I/σI | 12.5 (1.2)* |
| CC(1/2) | 0.998 (0.649)* |
| **Refinement** | |
| Resolution [Å] | 44.88 – 1.73 (1.76-1.73)* |
| No. of reflections (working/free) | 101840 / 5131 |
| R_cryst_ [%] | 18.1 |
| R_free_[%] | 21.5 |
| No. of atoms: | |
| Protein | 5801 |
| Water | 568 |
| Heterogens (inc ligand) | 60 |
| Deviation from ideal geometry: | |
| Bond lengths [Å] | 0.0149 |
| Bond angles [◦] | 2.13 |
| Ramachandran: | |
| Most favoured [%] | 96.9 |
| Allowed [%] | 3.1 |
| Disallowed [%] | 0.0 |

*Values in parenthesis refer to the highest resolution bin.

**Table S6. CNS Exposure in rats following peripheral dosing of small molecules and SDX-7320.** The test agents TNP-470 (dosing solution prepared in 10% ethanol/90% PBS at 6mg/ml), SDX-7539 (dosing solution prepared in PBS at 7.4 mg/mL) and SDX-7320 (dosing solution prepared in PBS at 40 mg/mL) were dosed intravenously at time zero (n=3/group). Animals were euthanized via CO_2_ inhalation 10 minutes post-dose (TNP-470, SDX-7539) or 120 minutes post-dose (SDX-7320) and blood was collected via cardiac puncture followed immediately by collection of cerebrospinal fluid (CSF). Blood was collected in to K_2_EDTA blood collection tubes for plasma separation; the CSF was collected from each rat then snap-frozen. Plasma and CSF samples were stored at -80^0^C until analysis.

| Test agent | Dose (mg/kg) | Analyte | Plasma, nM (SD) | CSF, nM (SD) | Plasma/CSF |
| --- | --- | --- | --- | --- | --- |
| TNP-470 | 30 | M-IV* | 1134 (453) | 460 (183) | 2.5 |
| SDX-7539 | 37 | SDX-7539 | 11,556 (3291) | 1,381 (1340) | 8.4 |
| SDX-7320 | 200 | SDX-7539 | 399 (12) | 27 (32) | 15 |

*active metabolite of TNP-470

**Table S7. Results of the rat FOB**. Male rats (approximately 8 weeks old) were individually housed 12 hours prior to dosing, to allow familiarity with the test chambers. SDX-7320 (prepared in 0.5% aqueous mannitol) was dosed once subcutaneously at either 3, 10 or 30 mg/kg and animals were observed twice/day for morbidity and mortality. The FOB was conducted at pre-dose, then 12- and 36-hours post-dose, by personnel who were blinded to the treatments each animal received. The only significant changes observed were in body weight (36-hour post-dose; 10 and 30 mg/kg groups).

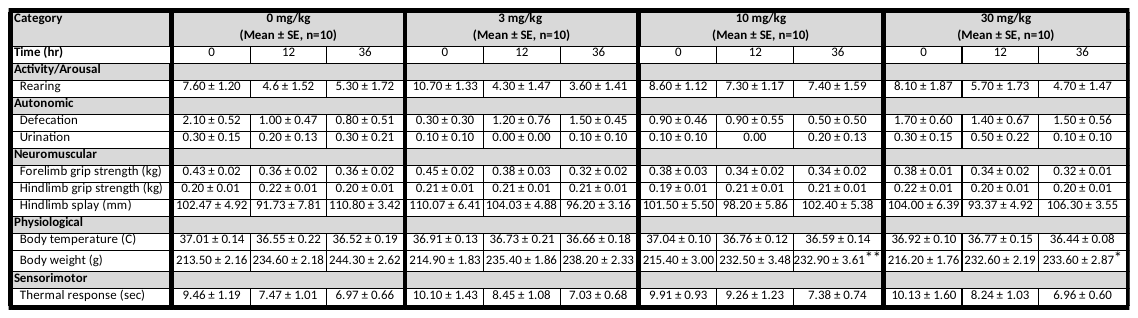

*p < 0.05, **p < 0.01 vs vehicle
